# An AGO2 adaptor expands the functional and evolutionary reach of microRNA targeting

**DOI:** 10.1101/2025.10.07.680889

**Authors:** Alex F. F. Crozier, Kunal M. Shah, Paul Grevitt, Anisha Thind, Eleni Maniati, Jun Wang, Kylie Shen, Diana Cox, Vinothini Rajeeve, Pedro Cutillas, Dimitris Lagos, Faraz Mardakheh, Sam Griffiths-Jones, Antonio Marco, Tyson V. Sharp

**Affiliations:** Centre for Cancer Cell and Molecular Biology, Barts Cancer Institute, Queen Mary University of London, London, UK; Centre for Cancer Biomarkers and Biotherapeutics, Barts Cancer Institute, Queen Mary University of London, London, UK; Eclipse BioInnovations, San Diego, CA, USA; Cell Signalling and Proteomics Laboratory, Centre for Cancer Evolution, Barts Cancer Institute, Queen Mary University of London, London, UK; York Biomedical Research Institute, University of York, York, UK; Department of Biochemistry, University of Oxford, South Parks Road, Oxford, UK; School of Biological Sciences, Faculty of Biology, Medicine and Health, University of Manchester, Manchester, UK; School of Life Sciences, University of Essex, Colchester, UK

## Abstract

Current models of microRNA (miRNA) silencing posit that RNA-sequence rules are sufficient for canonical targeting of mRNAs by Argonaute 2 (AGO2), the central protein of the miRNA-induced silencing complex (miRISC). Using chimeric eCLIP in CRISPR-edited LIMD1^+/+^, LIMD1^+/−^, and LIMD1^−/−^ human small airway epithelial cells (hSAECs), we reveal a transcriptome-wide dependency on LIMD1, an AGO2 adaptor, for effective miRNA targeting and repression. In LIMD1-deficient cells, miRNA loading is uncoupled from productive targeting: despite increased AGO2–miRNA interactions, complexes engage fewer transcripts and sites, reducing occupancy and more than halving both the breadth and depth of targeting. We also observe altered AGO2 positional footprints across targets in LIMD1-deficient cells. LIMD1 dependence is most pronounced at defined RNA contexts: weak (GC-poor) seed pairings, interactions involving evolutionarily young miRNAs or sites that nonetheless form thermodynamically stable duplexes, with these losses particularly enriched in coding sequences of rapidly evolving C_2_H_2_-zinc-finger genes. Even within canonical seed repertoires of individual AGO2–miRNAs, LIMD1 is most critical at poorly conserved sites, indicating that LIMD1 broadens miRNA regulation beyond ancient, deeply conserved targets. In culture, LIMD1 deficiency de-represses oncogenic proteins that, *in vivo*, inversely correlate with LIMD1 levels in normal lung and adenocarcinoma, where LIMD1 is characteristically reduced, and whose dysregulation predicts poor survival. Thus, LIMD1 emerges as a key determinant of miRISC architecture, targeting, and potency, challenging RNA-centric models of miRNA function and exemplifying how adaptor proteins diversify post-transcriptional regulation.

**Graphical Abstract:** LIMD1 defines the scope of miRNA-mediated targeting and repression

- AGO2-chimeric eCLIP in CRISPR-edited human small airway epithelial cells (hSAECs) shows LIMD1 is required for productive AGO2–miRNA engagement transcriptome-wide.
- LIMD1 deficiency reduces the AGO2–miRNA:targetome. Each AGO2–miRNA binds fewer targets, with lower occupancy per site and per transcript and fewer global silencing events.
- LIMD1 dependence is strongest for GC-poor seed-sites, less conserved miRNAs and sites, and thermodynamically stronger duplexes.
- Dosage-dependent effects of LIMD1 deficiency are broadly observed.
- Target-mRNA decay and translational repression are reduced in LIMD1-deficient hSAECs, increasing protein output.
- *In vivo*, LIMD1 LOH–associated deficiency is prevalent and typically clonal in NSCLC. LIMD1 expression inversely correlates with oncogene levels in normal lung and adenocarcinoma, and target dysregulation predicts poor survival.

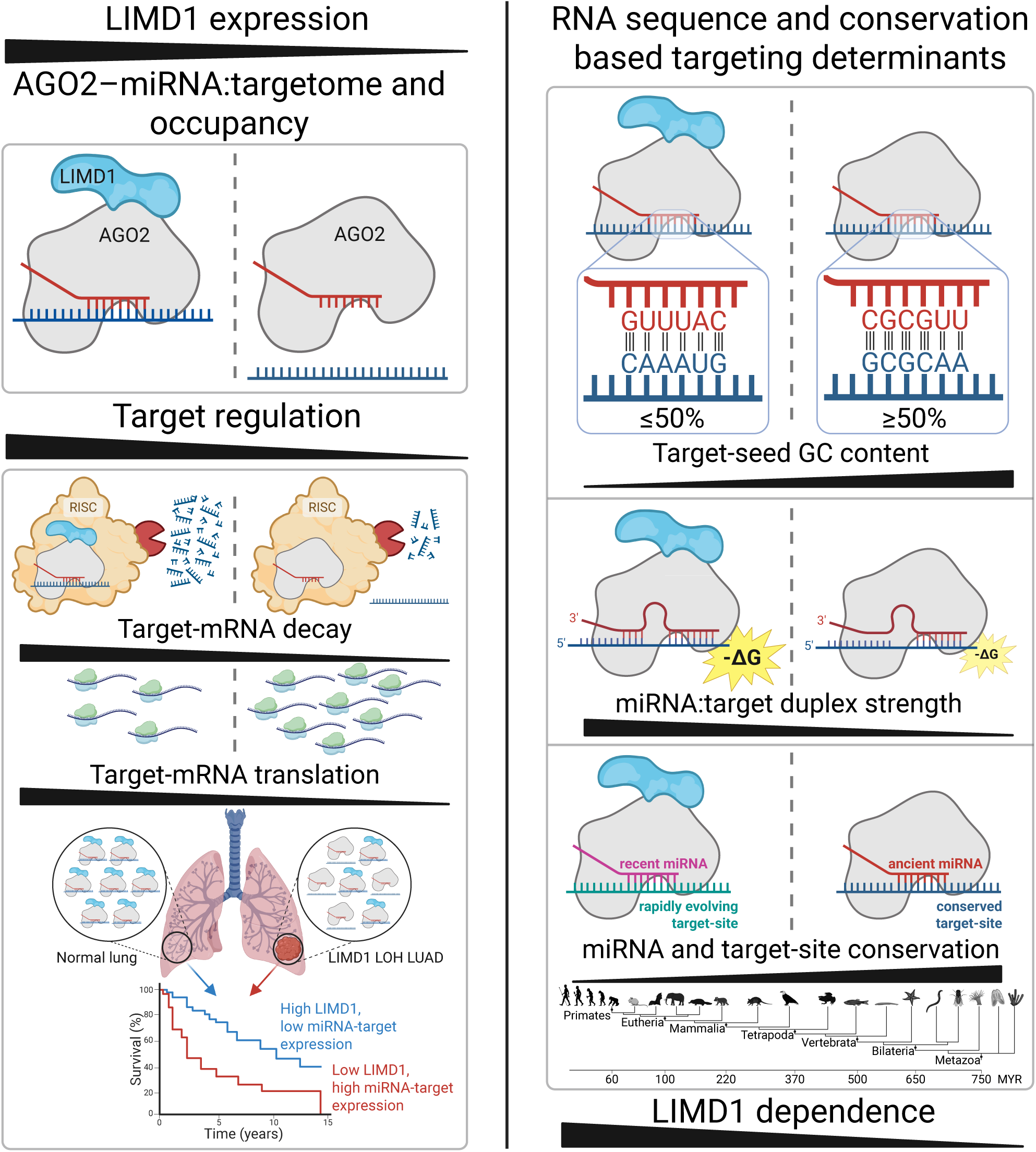

## Introduction

MicroRNAs (miRNAs) are small non-coding RNAs that repress gene expression post-transcriptionally by guiding Argonaute (AGO) proteins to target sites in mRNAs (*1*). Each miRNA can regulate many transcripts, and most human mRNAs carry conserved sites, together creating dense regulatory networks that shape development, homeostasis, and disease (*2*). Canonical targeting is dictated primarily by seed complementarity, with flanking sequence, supplementary pairing, and local structure modulating efficacy; non-canonical sites further broaden targeting (*1*, *3–8*). AGO–miRNA:target complexes recruit TNRC6 scaffolds and associated cofactors to form the miRNA-induced silencing complex (miRISC), which couples target recognition to translational repression and mRNA decay (*9–11*). Yet despite extensive study, the rules governing targeting efficacy remain incomplete, limiting mechanistic insight and therapeutic progress (*12*).

Although base-pairing explains site recognition, repression efficacy also reflects miRISC architecture (*13*, *14*). AGO–miRNAs are structurally dynamic, undergoing conformational remodelling upon target binding; post-translational modifications tune AGO interactions, TNRC6 multivalency stabilizes cooperative AGO binding to prolong target dwell time, and paralog choice adds specificity (*14–23*). These features support a model in which RNA base-pairing rules integrate with dynamic miRISC architecture to define repression specificity, strength, and kinetics (*13*, *15*, *24*, *25*). Yet most studies of targeting remain RNA-centric (*12*): despite recent advances, predictors of binding and repression remain unreliable (*8*), while evolutionary analyses emphasize miRNA and site conservation (*2*, *26*, *27*), leaving the role of miRISC adaptor proteins in targeting and regulatory evolution underexplored.

LIMD1, a metazoan adaptor-scaffold with tumour-suppressive activity in lung cancer (*28*–*30*), provides a rare mechanistically defined example. We previously mapped LIMD1–AGO2 and LIMD1–TNRC6A interactions, establishing LIMD1 as an adaptor/scaffold within miRISC that potentiates AGO2–TNRC6 binding and reporter silencing cells (*19*, *31*). Yet whether LIMD1 is globally required for AGO2–miRNA targeting, and what features determine such dependence, remain unknown.

Here, we take an integrated approach, combining AGO2-chimeric eCLIP (*32*) with transcriptome, proteome, motif, evolutionary, and thermodynamic analyses, in primary human small airway epithelial cells (hSAECs) across graded LIMD1 expression states. Our findings establish LIMD1 as a paradigm for how adaptor proteins shape AGO2–miRNA targeting, demonstrating that miRISC architecture is a central determinant of post-transcriptional gene regulation, with broad implications for physiology, disease, and evolution.

## Results

### LIMD1 potentiates AGO2–TNRC6A interactions and miRNA-reporter repression

Reduced LIMD1 expression, often via clonal loss of heterozygosity (LOH), is pervasive in non-small cell lung cancer (NSCLC) and predicts poor survival (*28*, *29*, *33*). We therefore used immortalised, non-transformed hSAECs, a diploid ATII-derived primary-line that shares lineage with lung adenocarcinoma (LUAD) (*34*, *35*), to study LIMD1’s role in miRNA silencing in a physiological and disease-relevant context.

CRISPR-Cas9 mediated editing generated three independent hSAEC clones per genotype: LIMD1 knockout (KO), heterozygous (Het, modelling LOH), and control (Ctrl) cells (***Fig. S1A–B***). Genome sequencing confirmed on-target editing with minimal off-target effects (***Table S1–3***). AGO2 protein was unchanged in KO and increased in Het, while TNRC6A was higher in both (***Fig. S1C***), excluding reduced AGO2 or TNRC6A protein abundance as the cause of any observed silencing defects.

Corroborating our previous study in HeLa cells (*19*), co-immunoprecipitation (co-IP) and *in situ* proximity ligation assays (PLA) in hSAECs showed that LIMD1 loss reduces AGO2– TNRC6A interaction (***Fig. S1D–G***) and derepresses miRNA reporters in a dose-dependent manner (***Fig. S1H***), establishing LIMD1 as the AGO2–TNRC6A scaffold for silencing in a non-tumour, diploid airway model.

### LIMD1 deficiency increases AGO2–miRNA interactions

To examine AGO2–RNA interactions across LIMD1 states, we performed AGO2 chimeric eCLIP (*32*) on our hSAEC panel (***Fig. S2A; Table S4–5***). The conventional (non-chimeric) eCLIP component of this method maps and thresholds AGO2-bound RNAs as reproducible peaks, quantifying AGO2–miRNA and AGO2–mRNA transcript association and peak enrichment.

Focusing first on reproducible AGO2–miRNA peaks, AGO2–miRNA binding increased dose-dependently with LIMD1 loss. LIMD1-deficient cells had an expanded AGO2-bound miRNA repertoire (Ctrl: 86; Het: 104; KO: 118), greater total AGO2–miRNA binding (+39% in Het, +88% in KO), and elevated AGO2 enrichment per matched peak and per miRNA (***Fig. 1A– F; S2B–E***).

**Figure 1.**
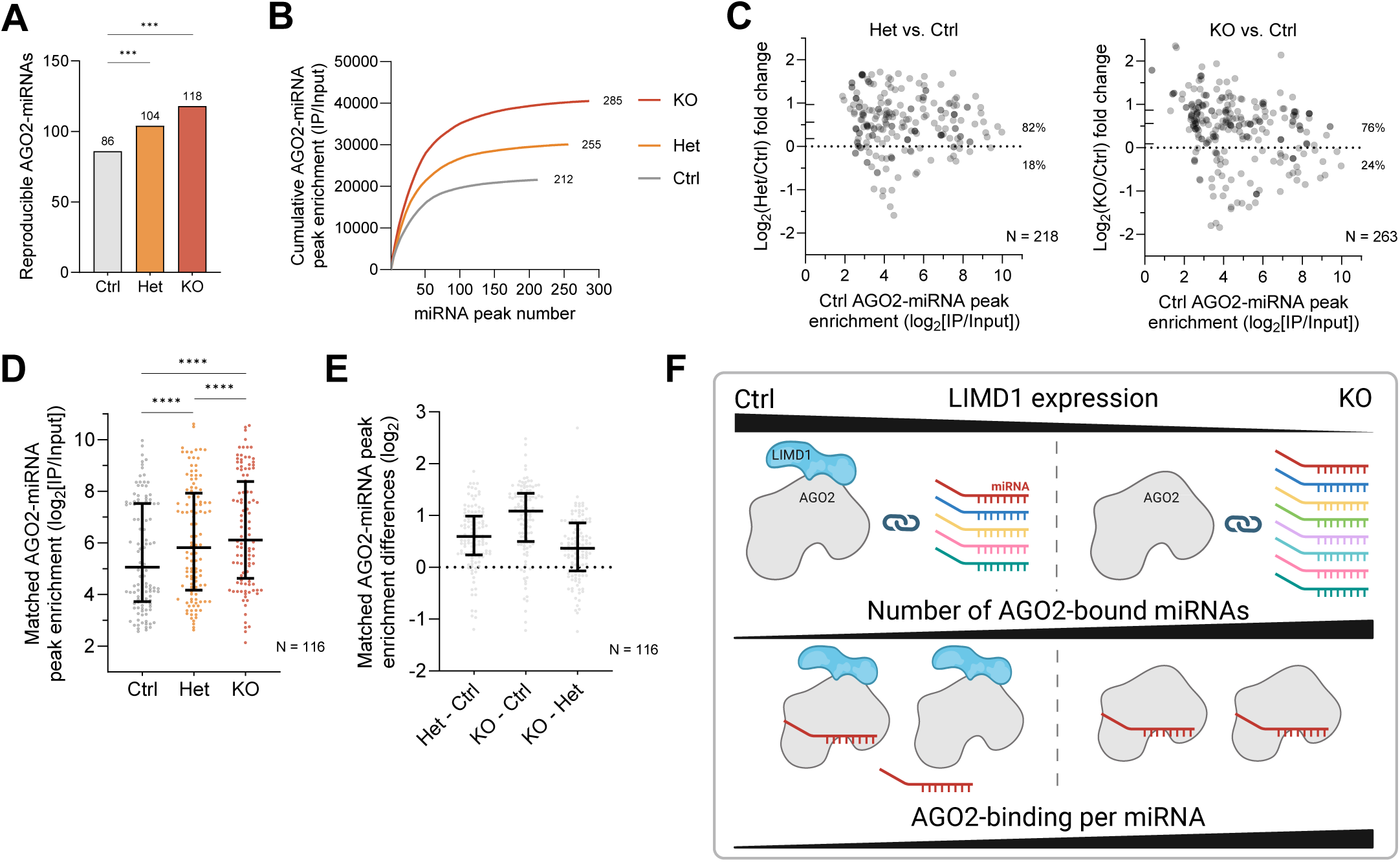
LIMD1 deficiency increases AGO2–miRNA interactions. AGO2-eCLIP analysis of reproducible miRNA peaks in hSAECs. **(A)** AGO2–miRNA repertoire expands with LIMD1 deficiency (pairwise proportion test with Benjamini–Hochberg correction; ***P<0.001). **(B)** Cumulative enrichment is higher in Het and KO cells, indicating dose-dependent increases in AGO2 loading. **(C)** Matched peak enrichment in Het and KO versus Ctrl. Each point represents a peak (x-axis: mean Ctrl enrichment; y-axis: fold change from DESeq2). Most peaks are enriched (percentages indicated); ticks to the right of the y-axis denote median and interquartile range. **(D, E)** Enrichment **(D)** and enrichment differences **(E)** for 116 peaks shared by Ctrl, Het, and KO, demonstrating a global, dose-dependent increase in AGO2–miRNA binding (Wilcoxon matched-pairs signed-rank test with multiple-comparisons correction; ****P<0.0001). **(F)** Schematic illustrating LIMD1 dose–dependent increases in the AGO2–miRNA repertoire and binding per miRNA in Het and KO cells versus Ctrl.

Although mature miRNAs tended to have higher expression in LIMD1-deficient cells, increased AGO2–miRNA association was independent of increased expression: miRNA abundance did not predict AGO2 enrichment and changes in expression and enrichment were negatively correlated (***Fig. S2F–L***). Thus, LIMD1 deficiency broadly increases AGO2– miRNA binding independent of increased miRNA expression, raising the question of whether this alters AGO2–mRNA interactions.

### AGO2–mRNA binding requires LIMD1

Despite increased AGO2–miRNA loading, analysis of AGO2–mRNA peaks revealed that LIMD1 loss reduced AGO2–mRNA binding. LIMD1-deficient cells had fewer AGO2- associated mRNAs (Ctrl: 912; Het: 523; KO: 659) and AGO2–mRNA peaks (5093; 2407; 3213), reduced total AGO2–mRNA binding, and lower median AGO2 enrichment per peak (−39% in Het, −51% in KO) (***Fig. 2A–E; S3A–D***). AGO2–mRNA binding losses were transcriptome-wide and evident across all mRNA regions (***Fig. 2F; S3C–H***).

**Figure 2.**
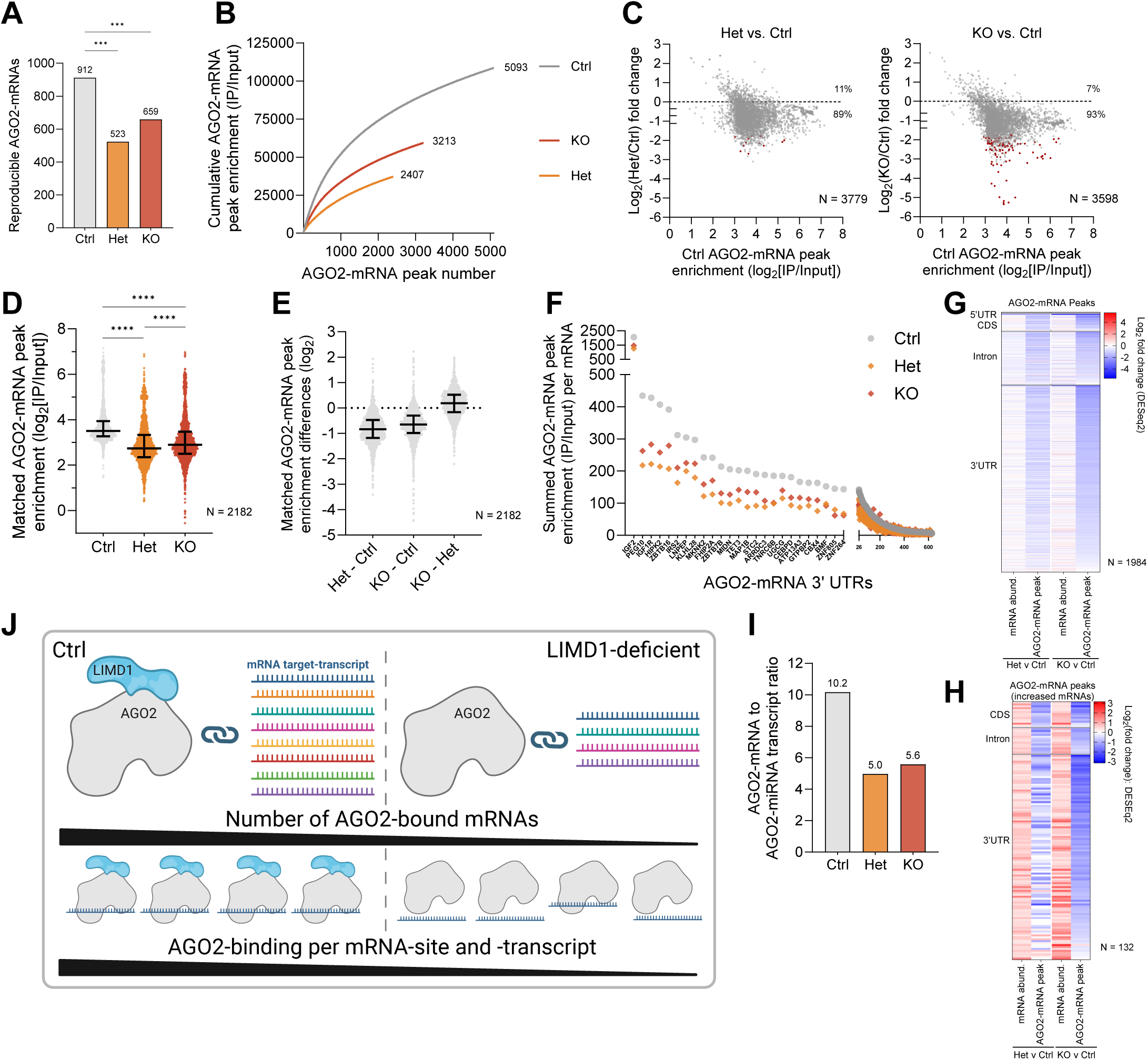
AGO2–mRNA binding requires LIMD1. AGO2-eCLIP analysis of reproducible mRNA peaks in hSAECs. **(A)** AGO2–mRNA repertoire contracts with LIMD1 deficiency (pairwise proportion test with Benjamini–Hochberg correction; ***P<0.001). **(B)** Reduced total AGO2–mRNA binding in LIMD1-deficient cells indicated by cumulative AGO2–mRNA enrichment. **(C)** Matched peak enrichment in Het and KO versus Ctrl. Each point represents a peak (x-axis: mean Ctrl enrichment; y-axis: fold change from DESeq2). Most peaks are de-enriched (percentages indicated); ticks to the right of the y-axis denote median and interquartile range; significant changes (|log₂FC|>1, q<0.1) are highlighted in red. **(D, E)** Enrichment **(D)** and enrichment differences **(E)** for 2,182 peaks common to Ctrl, Het, and KO, showing a global reduction in AGO2–mRNA binding (Wilcoxon matched-pairs signed-rank test with multiple-comparisons correction; ****P<0.0001). **(F)** Summed AGO2–mRNA enrichment across 3′ UTRs per transcript shows reduced binding in Het and KO versus Ctrl. **(G, H)** Heatmaps of log₂FC in mRNA abundance and AGO2–mRNA enrichment (IP/Input) for shared peaks, showing widespread loss of AGO2–mRNA binding **(G)** irrespective of transcript levels and **(H)** even when mRNAs are increased. Heatmaps are grouped by genomic feature and ordered by loss of binding in KO versus Ctrl. **(I)** Ratio of distinct AGO2-associated mRNAs to miRNAs is halved in LIMD1-deficient cells, indicating uncoupling of miRNA loading from productive mRNA targeting. **(F)** Schematic illustrating reduced AGO2–mRNA repertoire and binding per site and transcript in LIMD1-deficient cells compared with Ctrl.

These losses were independent of mRNA abundance, which showed no global decrease; correlations between mRNA expression and binding changes were weak, and peak enrichment declined even on transcripts with stable or increased abundance (***Fig. 2G–H; S3I–K***). Thus, independent of RNA abundance, LIMD1 deficiency uncouples AGO2–miRNA loading from productive AGO2–mRNA engagement, reducing the AGO2–mRNA to AGO2– miRNA ratio by half despite an expanded AGO2–miRNA repertoire (***Fig. 2I–J***).

### LIMD1 deficiency constrains AGO2–miRNA targeting breadth and depth

We next asked how LIMD1 deficiency alters AGO2–miRNA:target interactions, both transcriptome-wide and at the level of individual miRNAs, targets, and binding events. For this, we analysed AGO2 chimeric eCLIP data. Unlike conventional eCLIP, which measures RNA-peak enrichment, chimeric eCLIP captures ligated AGO2–miRNA:target complexes at nucleotide resolution (*32*). Each chimeric peak represents a unique, experimentally defined AGO2–miRNA:target-site interaction with its seed match, and each chimeric read corresponds to a single interaction event in cells.

LIMD1 deficiency resulted in a profound contraction of AGO2–miRNA:target interactions. Reproducible chimeric peaks fell by ∼60% (Ctrl: 1,661; Het: 667; KO: 652), with fewer target-bound AGO2–miRNAs (111; 84; 85) and a >50% decrease in miRNA-bound transcripts (734; 344; 331) (***Fig. 3A–C***). This was accompanied by a >50% decrease in total chimeric reads (quantifiable silencing events) and unique AGO2–miRNA:mRNA combinations (***Fig. 3D–E; S4A***). Thus, LIMD1 deficiency reduced both the global breadth and depth of AGO2–miRNA targeting, reflected in a contracted interaction repertoire, fewer target-engaged AGO2–miRNAs, a smaller pool of regulated mRNAs, and fewer total silencing events.

**Figure 3.**
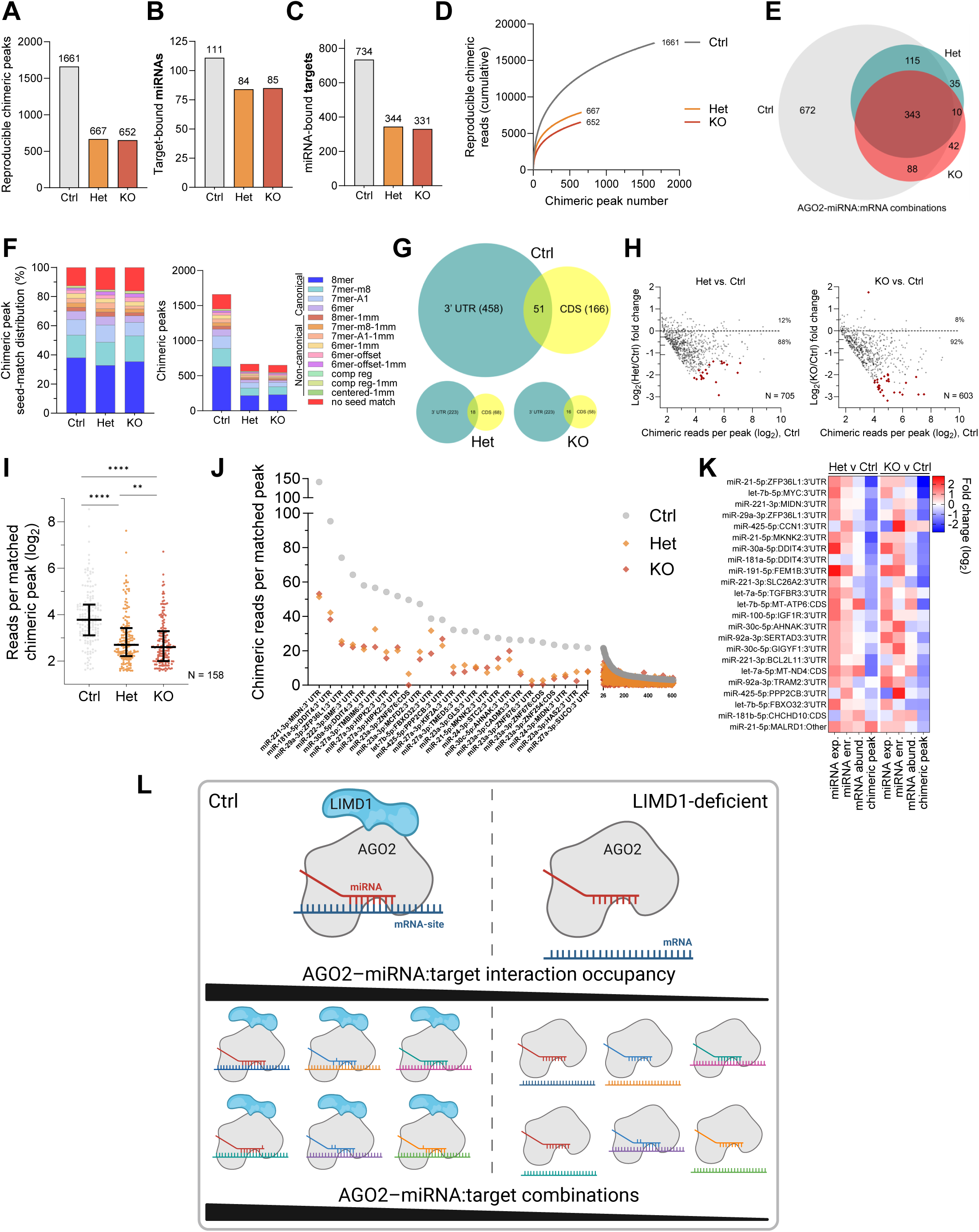
LIMD1 deficiency constrains global AGO2–miRNA targeting. AGO2-eCLIP chimeric analysis in hSAECs. **(A–C)** Numbers of reproducible chimeric peaks **(A)**, target-bound AGO2–miRNAs **(B)**, and AGO2–miRNA-bound target-transcripts **(C)** are reduced in LIMD1-deficient cells, revealing contraction of AGO2–miRNA targeting. **(D)** Cumulative chimeric read counts are lower in Het and KO, reflecting fewer AGO2– miRNA:target binding events. **(E)** Venn diagram of the AGO2–miRNA regulome (unique AGO2–miRNA:mRNA combinations) shows reduced regulatory breadth. **(F)** Seed-match distribution and number of chimeric peaks are reduced across all pairing classes (TargetScan criteria). **(G)** Proportional Venn diagrams of partially overlapping AGO2–miRNA targets in 3′ UTRs and CDSs indicate distinct regulatory subnetworks. **(H)** Matched chimeric peaks with ≥3 normalized reads in Ctrl. Each point represents a peak (x-axis: mean Ctrl reads; y-axis: fold change in Het or KO). Most peaks are de-enriched (percentages indicated); ticks right of the y-axis mark median and interquartile range; significant changes (|log₂FC|>1, *P*<0.05) are in red. **(I)** Normalized chimeric read counts for 158 peaks common to all groups (≥3 reads each), showing global reduction (Wilcoxon matched-pairs signed-rank test with multiple-comparisons correction; **P<0.01, ****P<0.0001). **(J)** Ranked chimeric reads per peak, with the 25 most abundant labelled by miRNA:mRNA:region. **(K)** Heatmap of log₂ fold changes in miRNA expression, AGO2–miRNA enrichment, mRNA abundance, and chimeric read counts for peaks with ≥20 reads in Ctrl and complete data, showing systematic loss of AGO2– miRNA:target-site occupancy in Het and KO despite increased miRNA expression and AGO2 loading. **(L)** Schematic illustrating reduced AGO2–miRNA:target interaction occupancy and repertoire in LIMD1-deficient cells.

All seed-match classes were affected, with similar relative distributions indicating ∼uniform effects across canonical and noncanonical pairings (***Fig. 3F; S4B–C***). Losses spanned all transcript regions, but were disproportionate in coding sequences (CDSs) (***Fig. S4D–F***); the minimal overlap between CDS and 3′UTR targets reveals distinct regional subnetworks of regulation, both dependent on LIMD1 (***Fig. 3G***). Across matched chimeras, median read counts fell by 36% in Het and 51% in KO (***Fig. 3H–I; S4G–H***), a trend persisting among the most frequent interactions and despite increased miRNA expression and AGO2 loading (***Fig. 3J–L; S4I***).

At the level of individual miRNAs, targeting capacity contracted sharply: median chimeric reads per miRNA halved, and each AGO2–miRNA bound fewer sites and transcripts (***Fig. 4A–E; S5A–I***). For instance, miR-23a-3p and miR-27a-3p targeted >90 mRNAs in Ctrl but <30 in Het or KO. Accordingly, binding events per mRNA declined by a median of 48% in Het and 63% in KO, with fewer distinct AGO2–miRNAs per transcript across CDS and 3′UTRs (***Fig. 4F–I; S5J–P***). Representative networks highlight this collapse, with prevalent AGO2–miRNAs (miR-23a-3p, let-7a-5p, and miR-27a-3p) engaging fewer targets and at reduced frequencies per interaction and per mRNA 3′UTR (***Fig. 4J–M***). Together, these system-wide and miRNA-specific data quantify the reduced breadth, occupancy, and depth of the AGO2–miRNA targeting network in LIMD1-deficient cells, establishing a dose-dependent requirement for LIMD1 in miRNA targeting (***Fig. 4N***).

**Figure 4.**
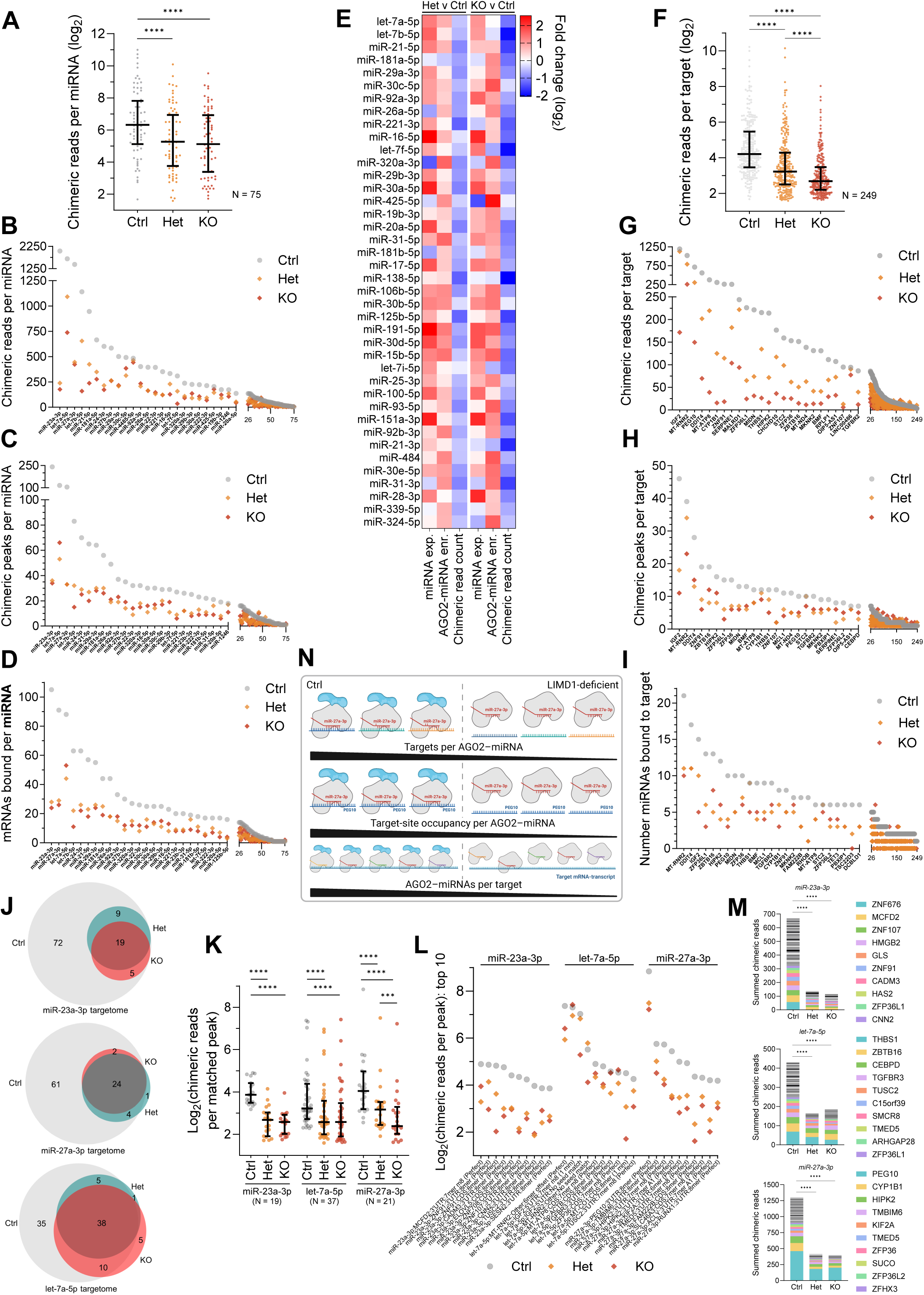
LIMD1 deficiency constrains breadth and depth of targeting by individual AGO2–miRNAs. Per-miRNA and per-target AGO2-eCLIP chimeric analysis in hSAECs. **(A)** Chimeric reads per AGO2–miRNA across reproducibly detected miRNAs show dose-dependent reduction in LIMD1-deficient cells (Wilcoxon matched-pairs signed-rank test; ****P<0.0001). **(B–D)** Expanded views of the top 25 (left) and next 50 (right) AGO2–miRNAs show reduced chimeric reads **(B)**, peaks **(C)**, and target transcripts **(D)** per miRNA, ordered by counts in Ctrl. **(E)** Fold change (log₂) in mature miRNA expression, AGO2–miRNA enrichment, and chimeric reads per miRNA (restricted to miRNAs with complete data), showing widespread loss of total target binding per AGO2–miRNA despite increased expression and AGO2 loading. **(F)** Summed chimeric reads per reproducibly detected AGO2–miRNA:*target* show dose-dependent reduction in LIMD1-deficient cells (Wilcoxon matched-pairs signed-rank test; ****P<0.0001). **(G–I)** Expanded views of the top 25 (left) and next 224 (right) target transcripts show reduced chimeric reads **(G)**, peaks **(H)**, and bound miRNAs **(I)** per transcript, ordered by counts in Ctrl. **(J)** Proportional Venn diagrams of the AGO2–miRNA targetome show reduced targeting breadth for the top three Ctrl miRNAs (miR-23a-3p, let-7a-5p, miR-27a-3p). **(K, L)** Chimeric reads per matched peak (≥3 reads in each) for these miRNAs show reduced occupancy per site in LIMD1-deficient cells, extending beyond the targetome losses in (J). **(L)** Read counts for the top 10 matched interactions are labelled by miRNA:target:region:seed-pairing. **(M)** Summed chimeric reads per 3′UTR for transcripts targeted by miR-23a-3p, miR-27a-3p, and let-7a-5p show reduced total reads, target number, and reads per 3′UTR in LIMD1-deficient cells, highlighting reduced targeting depth. Top 10 targets per AGO2–miRNA in Ctrl are indicated by colour in each group. **(N)** Schematic illustrating reduced targets and target-site occupancy per AGO2–miRNA, and reduced AGO2–miRNAs per target-transcript in LIMD1-deficient cells.

### LIMD1 shapes AGO2–miRNA targeting landscapes by governing motif preferences and positional footprints

To resolve how LIMD1 shapes AGO2–miRNA targeting specificity, we applied positionally enriched k-mer analysis (PEKA) (*36*) to our AGO2-chimeric eCLIP data. PEKA identifies and quantifies enriched sequence motifs and their positional distribution relative to thresholded AGO2-crosslink sites, enabling global and nucleotide-level analysis of target-binding preferences.

LIMD1 deficiency reshaped the target-motif landscape. Of the top 40 k-mers in Ctrl cells, only 9 and 6 were retained in Het and KO, respectively, with 15 shared between Het and KO (***Fig. 5A–C; S6A***), indicating progressive and wholesale reconfiguration of the sequence repertoire bound by AGO2–miRNAs. Notably, prevailing motifs in Het and KO were consistently more GC-rich (***Fig. 5D***), prompting us to test whether GC-poor canonical seed interactions were disproportionately affected by LIMD1 deficiency, which indeed they were (***Fig. 5E–G; S6B***). Since higher GC content confers greater base-pairing stability through additional Watson-Crick bonds (*37*), this suggests that weaker, GC-poor seed:target chimeras have heightened LIMD1 dependence.

**Figure 5.**
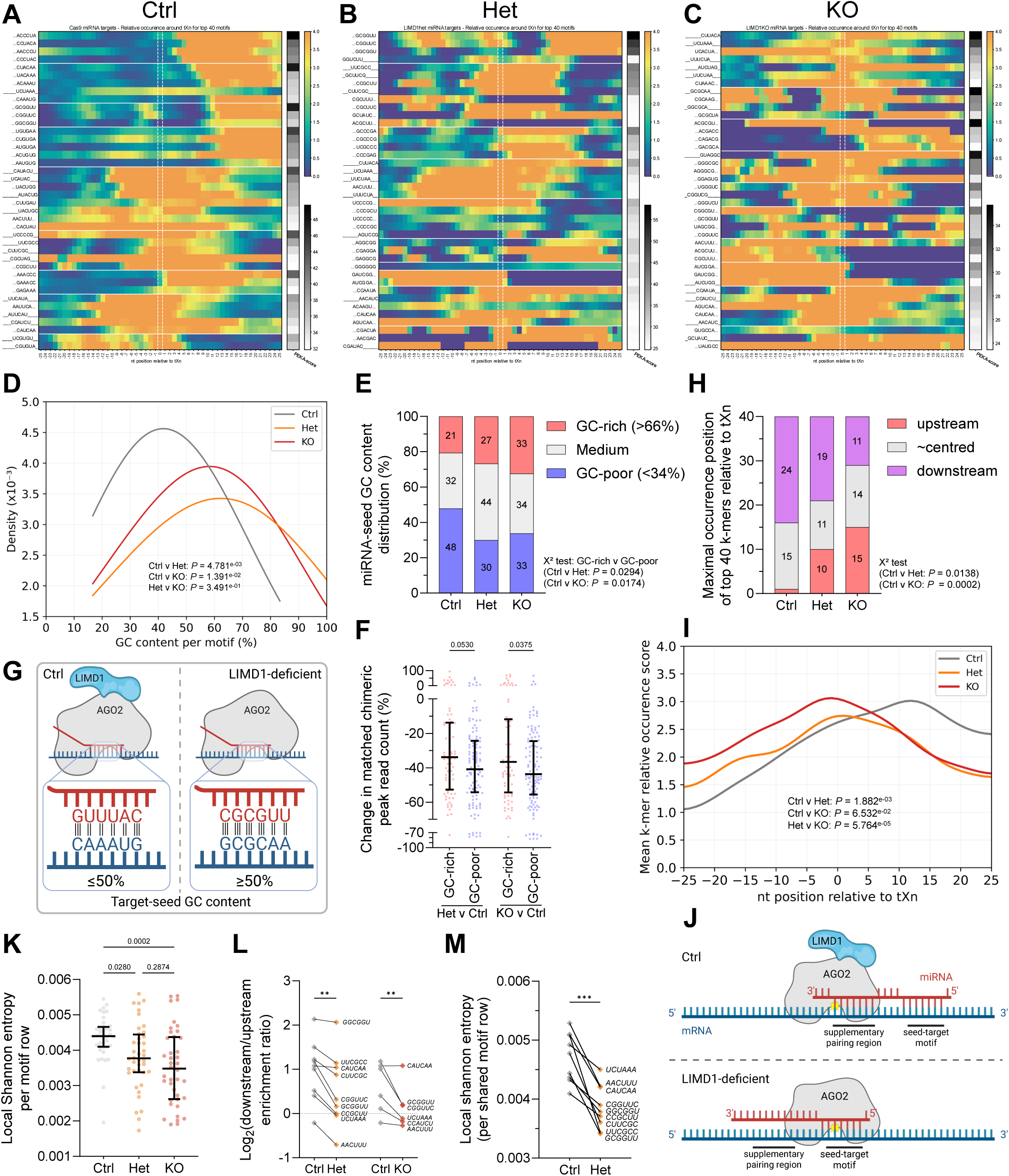
LIMD1 shapes AGO2–miRNA targeting landscapes by governing motif preferences and positional footprints. **(A–C)** Positionally enriched k-mer analysis (PEKA) of the top 40 target motifs from reproducible chimeric peaks in Ctrl **(A)**, Het **(B)**, and KO **(C)**. Heatmaps depict motif relative occurrence (RtXn), defined as fold enrichment of k-mers near thresholded crosslink (tXn) sites relative to distal background. PEKA scores reflect enrichment strength and positional consistency. K-mers are clustered by sequence similarity and aligned by maximal RtXn; dotted lines indicate cases where maximal enrichment occurs >3 nt from the crosslink site. LIMD1 deficiency alters motif identity, positional distribution, and binding footprints. Enlarged versions of heatmaps are provided in Fig. S9A–C for clarity. **(D)** GC content of the 40 most prevalent motifs, showing a shift toward GC-rich motifs in LIMD1-deficient cells (kernel density, Gaussian σ = 2.5; Kruskal–Wallis with multiplicity correction). **(E)** Distribution of GC-rich (>66% GC content), medium, and GC-poor (<34%) canonical seed types in Ctrl, Het, and KO, showing a dose-dependent increase in GC-rich seeds among remaining interactions in LIMD1-deficient cells (χ² test). **(F)** Change in matched chimeric peak read counts (%) for GC-rich (>66%) and GC-poor (<34%) canonical seeds in Het and KO versus Ctrl, showing greater loss of GC-poor seeds (Kruskal–Wallis test). **(G)** Schematic illustrating the typical GC content of canonical seed-interactions in Ctrl and LIMD1-deficient cells. **(H)** Distribution of maximal motif enrichment positions relative to crosslink sites, showing loss of the downstream (3′) bias in Ctrl and a dose-dependent upstream (5′) shift in LIMD1-deficient cells (χ² test). **(I)** Mean relative enrichment per nucleotide position for the 40 most prevalent motifs, showing loss of downstream bias and a shift toward upstream binding (kernel density, Gaussian σ = 3.33; multiplicity-corrected Mann–Whitney U test). **(J)** Interpretive schematic, aligned to nt positions in (I), illustrating the shift in peak motif enrichment relative to the AGO2 crosslink site (yellow starburst denotes the UV-induced crosslink site between AGO and target RNA [tXn]): in Ctrl, motif enrichment peaks ∼12 nt downstream of tXn, whereas in Het and KO it is crosslink-proximal (–2 nt). This 14-nt shift reflects re-positioning of AGO2–target contact sites from upstream of the seed in Ctrl to seed-proximal in Het and KO, consistent with loss of upstream contacts mediating supplementary pairing. **(K)** Local Shannon entropy per motif, quantifying variability in AGO2–miRNA binding across nucleotides. Higher entropy reflects broader, flexible footprints; lower entropy indicates punctate enrichment. LIMD1-deficient cells show reduced entropy per motif compared with Ctrl (multiplicity-corrected ANOVA). **(L, M)** Motifs shared between Ctrl and Het (n=9) or Ctrl and KO (n=6); sequences indicated. **(L)** Relative downstream/upstream enrichment ratio (log₂) shows a shift from downstream (3′) to upstream (5′) bias in LIMD1-deficient cells. **(M)** Local Shannon entropy per motif is reduced, indicating loss of footprint flexibility. Together these results show that LIMD1 governs AGO2 positional preference and binding flexibility even for identical motifs (corrected matched one-way ANOVA; **P<0.01, ***P<0.001).

LIMD1 deficiency also altered AGO2’s positional footprint on target motifs. In Ctrl cells, motif enrichment typically peaked >3 nt downstream of the AGO2 crosslink site, with a positive downstream-to-upstream enrichment ratio (***Fig. 5H–J; S6C***), a pattern expected if the predominant AGO2–target contact is upstream of the target-seed match, where it can mediate miRNA 3′ supplementary pairing (*3*, *7*, *38*). In LIMD1-deficient cells, this downstream bias was lost and enrichment shifted toward seed-proximal positions, consistent with disrupted supplementary contacts. LIMD1 deficiency was also associated with disruption of intermediate-strength contacts, with more zero-occurrence and fewer moderately enriched nucleotide-sites (***Fig. 5A–C; S6D***). Supporting this, local Shannon entropy per motif (which measures variability in enrichment across adjacent nucleotides) was higher in Ctrl, reflecting smoother binding-enrichment gradients and therefore broader, more flexible footprints (***Fig. 5K***). By contrast, Het and KO showed reduced entropy, corresponding to a more punctate, disordered binding profile. Critically, comparisons of shared motifs confirmed these shifts: in LIMD1-deficient cells, motif enrichment shifted 5′ relative to the crosslink site and footprint entropy was reduced (***Fig. 5L–M; S6E–H***), showing that LIMD1 governs AGO2’s positional preference and footprint flexibility at identical sequence motifs.

Thus, LIMD1 specifies both which motifs are engaged and AGO2’s positional footprint on target mRNAs. In its absence, AGO2–miRNA targeting contracts to GC-rich motifs with upstream-shifted, punctate footprints, causing disproportionate loss of weaker seed interactions and erosion of regulatory breadth and flexibility.

### LIMD1 deficiency de-represses miRNA targets in hSAECs and LUAD

Loss of AGO2–miRNA:target interactions caused widespread de-repression of downstream targets. Quantitative proteomics of our hSAEC panel revealed increased protein abundance for transcripts with 3′UTR chimeric sites, with levels scaling both with the degree of LIMD1 loss and the extent of chimeric peak de-enrichment (***Fig. 6A; S7A***). Parametric Analysis of Gene Expression (PAGE) (*39*) confirmed coordinated shifts: AGO2–miRNA targets defined by conventional or chimeric eCLIP were elevated, with the most de-enriched 3′UTR chimeric sites yielding the highest Z-scores (***Fig. S7B***). PAGE analysis further showed that LIMD1 deficiency de-repressed entire AGO2–miRNA targetomes: across 11 chimeric-CLIP–defined sets, five were significantly increased in Het cells (miR-27a-3p, miR-26a-5p, miR-181a-5p, miR-24-3p, miR-27b-3p) and seven in KO cells (additionally let-7a-5p and let-7b-5p) (***Fig. 6B; S7C–D***). Consistently, de-enriched chimeric targets (p < 0.1) and entire AGO2–miRNA targetomes showed elevated protein levels despite increased miRNA abundance and AGO2 association (***Fig. 6C–E; S7E***). Mechanistically, both stable and upregulated transcripts showed accumulated protein, indicating loss of translational repression as well as mRNA decay (***Fig. 6F; S7F***).

**Figure 6.**
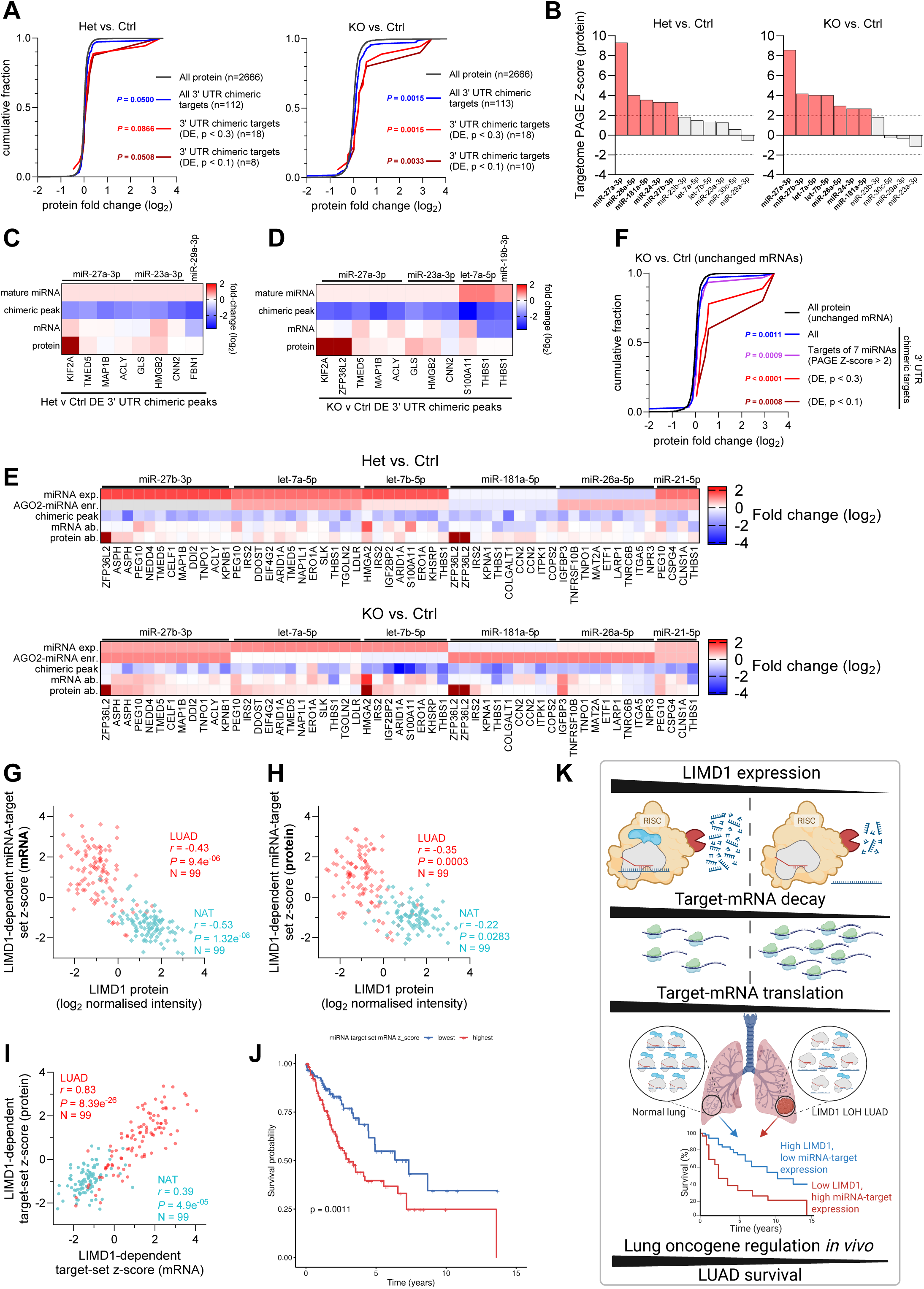
LIMD1 deficiency de-represses miRNA targets in hSAECs and LUAD. **(A)** Cumulative distribution of protein fold changes (log₂) in Het or KO versus Ctrl, measured by MS from six independent replicates of Ctrl C4, Het C6, and KO C2 hSAECs. Proteins with 3′UTR chimeric peaks (blue) show increased abundance relative to all proteins (black), strongest for the most de-enriched peaks (FDR<0.3, red; FDR<0.1, dark red; multiplicity-corrected Kruskal–Wallis test). **(B)** PAGE Z-scores of AGO2–miRNA targetomes show increased protein levels of chimeric targets; significant increases (Z >1.96 ≡ *P*<0.05) are highlighted in red. **(C–E)** Heatmaps of log₂ fold changes in miRNA expression, AGO2–miRNA enrichment, chimeric reads, mRNA abundance, and protein abundance (restricted to targets with complete data, including protein). (C, D) De-enriched 3′UTR chimeric targets (FDR<0.1) in Het (C) and KO (D). (E) Full 3′UTR chimeric targetomes of selected miRNAs, showing consistent patterns across datasets (gray = missing data). Colour scale capped to enhance visibility of sub-maximal protein increases; log₂ change >3 shown as dark red. **(F)** Cumulative distribution of protein fold changes in KO versus Ctrl for proteins with unchanged mRNA abundance (0.8–1.2-fold). Protein levels are increased for 3′UTR chimeric targets (blue), targets of AGO2–miRNAs with PAGE Z-scores >2 (purple), and de-enriched chimeric targets (red/dark red), consistent with translational derepression independent of mRNA decay (multiplicity-corrected Kruskal–Wallis test). **(G, H)** In CPTAC LUAD samples, LIMD1 protein levels inversely correlate (Pearson) with target-set z-scores at mRNA **(G)** and protein **(H)** levels for five de-enriched 3′UTR targets upregulated in LIMD1-deficient hSAECs. **(I)** Correlation between target-set mRNA and protein z-scores is stronger in LUAD than NAT, consistent with loss of translational control in tumours. **(J)** Kaplan–Meier survival analysis of LUAD patients (TCGA-LUAD) stratified by target-set mRNA signature: patients in the highest quartile (red) had poorer survival than those in the lowest quartile (blue; log-rank *P* = 0.0011; n=129/group). **(K)** Schematic illustrating LIMD1-dependent control of target mRNA decay and translation; LIMD1 protein inversely correlates with *in vivo* target-oncogene expression; high target-oncogene (and/or low LIMD1) expression associates with poorer LUAD survival.

We next tested whether these effects extended beyond CRISPR-hSAECs. A five-gene target set (KIF2A, VDAC1, PPIF, HMGB2, IGFBP3), defined as LIMD1-dependent from multi-omic analyses within our hSAEC panel and subsequently tested in public lung datasets, inversely correlated with LIMD1 protein in LUAD tumours and matched normal adjacent tissue (NAT) (***Fig. 6G–H; S7G–J***). In tumours, both mRNA and protein levels of this set were elevated compared with NAT, and mRNA–protein correlation was stronger (r = 0.83 vs 0.39) (***Fig. 6I; S7H–J***), consistent with loss of translational control (*40*, *41*). Clinically, high target-set expression predicted poor survival and remained significant in multivariate models, with a LIMD1–target set composite score achieving even stronger stratification (***Fig. 6J; S7K–O***), linking LIMD1 deficiency, associated with clonal *LIMD1* LOH (*29*), with defective miRNA-mediated repression and adverse outcome in LUAD (***Fig. 6K***).

### LIMD1 augments evolutionarily young yet thermodynamically strong interactions, enriched in C_2_H_2_-zinc-finger genes

LIMD1 is a metazoan-specific adaptor protein within the Zyxin family, a lineage that diversified with the emergence of multicellular complexity (*30*). Given the profound disruption to AGO2–miRNA targeting in LIMD1-deficient cells, we asked whether LIMD1 preferentially supports recently evolved regulatory circuits, thereby contributing to the evolutionary expansion of miRNA networks.

We first tested whether AGO2–miRNAs that retained target-binding in LIMD1 KO (≥1 reproducible chimeric peak) were more conserved than those that lost target-binding entirely (no reproducible peaks). Indeed, those retaining target-binding traced mainly to bilaterian seed families and to loci arising from vertebrate whole-genome duplications, whereas those that lost binding were enriched for mammalian-specific seed families and loci (***Fig. 7A–C; S8A***) (*27*). Although older miRNAs were modestly more abundant, logistic regression showed that evolutionary conservation independently predicted retention of target-binding in KO (***Fig. S8B–C***). Moreover, LIMD1-dependent interactions, identified as 3′UTR chimeric peaks depleted in KO, were disproportionately associated with younger miRNAs (***Fig. 7D***). Together, these findings indicate that LIMD1 preferentially supports the activity of evolutionarily young miRNAs, thereby extending the breadth of AGO2–miRNA regulatory networks.

**Figure 7.**
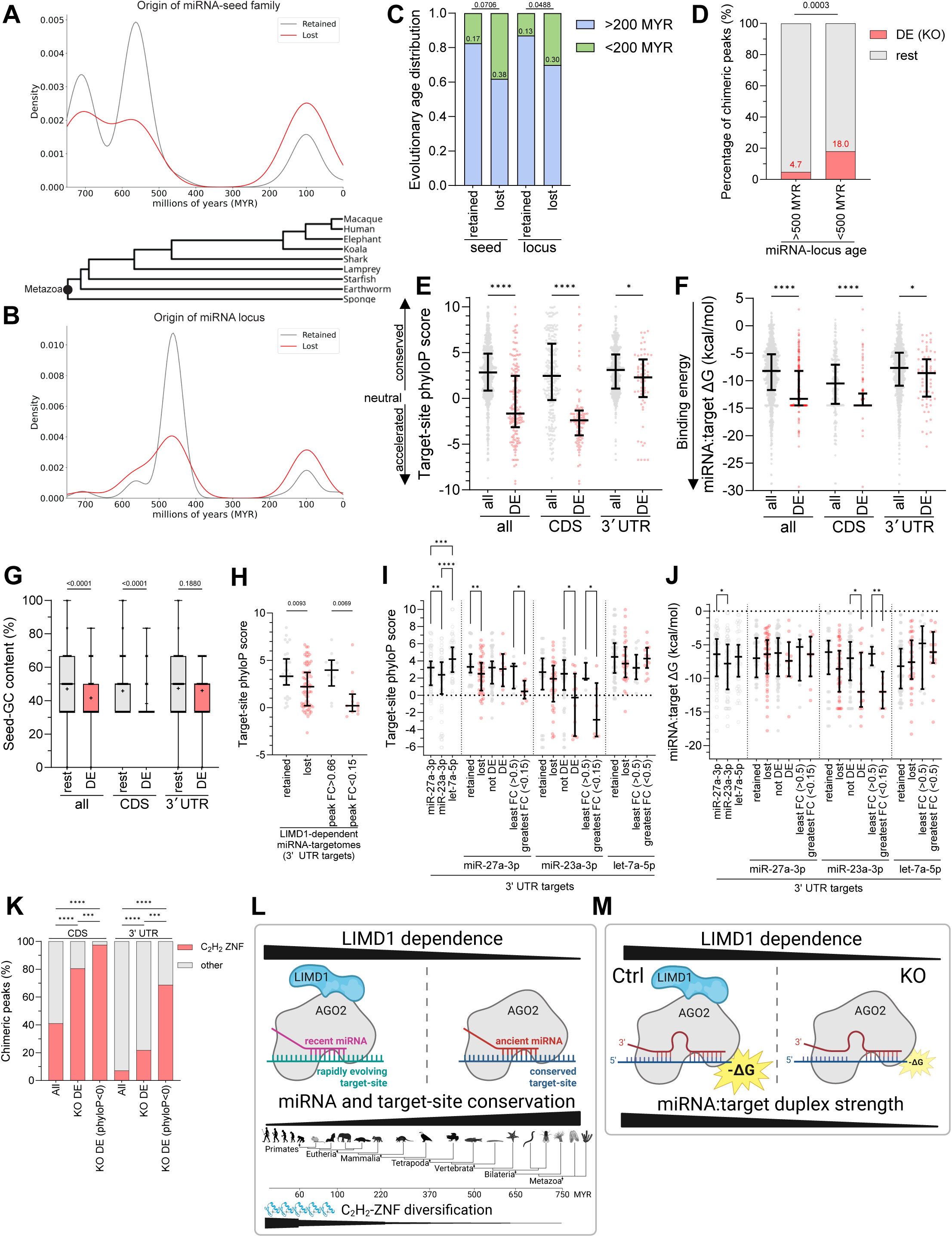
LIMD1 augments evolutionarily young yet thermodynamically strong interactions. **(A, B)** Kernel density distributions of evolutionary ages for miRNA seed families **(A)** and loci **(B)** reproducibly detected in chimeric peaks (“target-engaged miRNAs”). Families and loci which *retain* target-binding in KO (gray) trace to older origins, whereas those which *lost* target-binding in KO (red) skew toward mammalian-specific lineages. A phylogenetic tree provides context. **(C)** A higher proportion of younger (<200 MYR) than older (>200 MYR) miRNAs lose target binding in KO, indicating LIMD1 dependence (Fisher’s exact test). **(D)** LIMD1-dependent AGO2–miRNA:target interactions (depleted [DE] in KO, *P*<0.3) are disproportionately associated with younger miRNAs (<500 MYR; Fisher’s exact test). **(E, F, G)** LIMD1-dependent canonical interactions (DE peaks in KO, P<0.3) show lower site conservation (E), greater duplex stability (F), and lower seed-GC content (G) than overall CDS/3′UTR targets (multiplicity-corrected Kruskal–Wallis test). PhyloP scores >0 indicate increasing conservation, whereas <0 indicates accelerated evolution. Duplex stability was assessed by minimum free energy (ΔG), where increasingly negative values reflect stronger base pairing. Together these data indicate that the most LIMD1-dependent interactions are less conserved but thermodynamically stable. **(H)** For LIMD1-dependent miRNAs (miR-27a-3p, miR-27b-3p, miR-26a-5p, miR-24-3p, miR-181a-5p; Fig. 6B), sites lost or with high relative loss (FC<0.15) are less conserved than retained sites or those with minimal loss (FC>0.66; multiplicity-corrected Kruskal–Wallis test). **(I, J)** Canonical 3′UTR peaks of miR-27a-3p, miR-23a-3p, and let-7a-5p stratified by LIMD1 dependence: retained vs lost in KO, or by relative depletion (FC>0.5 vs <0.15 or <0.5). LIMD1-dependent sites for miR-27a-3p and miR-23a-3p have significantly lower phyloP scores, and for miR-23a-3p also more stable duplexes (more negative ΔG; right). Significance from multiplicity-corrected Kruskal–Wallis test; only *P*<0.05 shown. **(K)** LIMD1-dependent target sites (DE in KO, *P*<0.3) are strongly enriched for C_2_H_2_ zinc-finger (ZNF) genes, especially in CDS (87/108, 81%). Enrichment is even greater among sites with phyloP<0 (accelerated evolution; 79/81, 98% CDS; 11/16, 69% 3′UTR). Categories per HGNC; χ² test (***P<0.001, ****P<0.0001). **(L–M) (L)** Schematic illustrating that the most LIMD1-dependent interactions involve recent miRNAs or less conserved target sites, with an evolutionary tree, timeline and C_2_H_2_-ZNF diversification indicated for context. **(M)** LIMD1-dependent interactions also form stronger AGO2–miRNA:target duplexes (more negative ΔG).

We next analysed canonical interactions using phyloP conservation scores at target-sites (*42*) and predicted hybridization energies of 22-nt miRNA:target duplexes (*43*), stratified by target-region and LIMD1 dependence (***Fig. S8D–F***). LIMD1 dependence (–log₁₀ p-value) correlated with lower site conservation, reduced seed GC content, and greater duplex stability, largely driven by CDS interactions (***Fig. S8E*)**. Consistently, LIMD1-dependent canonical interactions were enriched at less conserved, lower-seed-GC target-sites that nonetheless formed stronger duplexes (more negative ΔG) (***Fig. 7E–G***), a pattern persisting even in transcripts with unchanged or increased abundance (***Fig. S8G***). Within 3′UTR targetomes of LIMD1-dependent miRNAs, the most depleted sites were less conserved (***Fig. 7H; S8H***). These trends extended to individual repertoires such as miR-27a-3p and miR-23a-3p, but not the more ancient let-7a-5p: LIMD1 dependence was greatest at less conserved sites forming more stable duplexes, showing canonical seed pairing alone is insufficient and that AGO2–miRNA engagement at these sites requires LIMD1 (***Fig. 7I–J***).

The most LIMD1-dependent sites were predominantly low-conservation, highly stable 8mers concentrated in C_2_H_2_-zinc-finger (C_2_H_2_-ZNF) transcripts (***Fig. 7K; S8I–L***), a rapidly evolving transcription factor family with central roles in DNA binding and regulation that underwent major lineage-specific expansions, especially pronounced in primates (*44–46*). Many of these sites exhibited negative phyloP scores, consistent with accelerated evolution, an anticipated feature of regulatory elements within this dynamically evolving gene family, which are prone to lineage-specific and clade-restricted diversification (*44*, *45*).

Together, these findings show that, beyond its broad role in augmenting miRNA-silencing, LIMD1 dependency is especially pronounced for interactions involving younger miRNAs or less conserved target-sites that form thermodynamically stable duplexes, particularly within CDS and C_2_H_2_-ZNF transcripts (***Fig. 7L–M***).

## Discussion

Our findings establish LIMD1 as a critical miRISC adaptor-scaffold that augments AGO2– miRNA binding and repression of targets across the transcriptome. In LIMD1-deficient cells, AGO2 loads more miRNAs yet engages far fewer transcripts and sites: individual targetome analyses showed each AGO2–miRNA bound fewer targets with reduced occupancy at remaining sites, while site-level analyses revealed fewer AGO2–miRNAs engaged per transcript and reduced binding across entire mRNAs. This uncoupling of AGO2–miRNA loading from productive targeting spans canonical and non-canonical sites, producing widespread protein de-repression that scales with LIMD1 loss and reflects impaired translational repression and mRNA decay. Integration of LIMD1 into miRISC therefore ensures AGO2–miRNA loading is coupled to effective post-transcriptional silencing.

These findings demonstrate that miRISC architecture, beyond miRNA abundance, AGO2 loading, or sequence features such as seed complementarity (*1*, *8*), determines AGO2– miRNA target engagement transcriptome-wide. LIMD1 imposes graded dependence, even within a single canonical seed repertoire; while base-pairing defines potential targets, adaptor incorporation into miRISC is required for stable engagement. Thus, LIMD1 modifies the outcome of sequence-defined interactions that would otherwise be less frequent, less stable, or absent. Together, these results extend RNA-centric models of miRNA function to include miRISC scaffold composition as a central determinant of targeting efficacy. Accordingly, predictive models of miRNA function and small-RNA therapeutic strategies must move beyond sequence features to account for miRISC architecture (*8*, *47*, *48*).

Mechanistically, LIMD1 acts as a miRISC scaffold that broadens AGO2’s targetome and sustains AGO2–miRNA:target interactions across both high- and low-occupancy sites, which collapse in its absence. This confinement of AGO2–miRNAs to a narrower set of binding- contexts may render kinetically challenging sites inaccessible without LIMD1. Indeed, GC-poor seed-motifs are disproportionately depleted, with AGO2–miRNAs contracting onto GC-rich, higher-affinity seeds in LIMD1’s absence (*37*). Paradoxically, the canonical interactions most dependent on LIMD1 are also predicted to form the most stable miRNA:mRNA duplexes, underscoring that thermodynamic favourability alone is insufficient for engagement. LIMD1 deficiency also produces upstream-shifted, rigid AGO2–target footprints at target-motifs, consistent with a shift from broad AGO-nucleotide sampling and 3′ supplementary bias to more seed-proximal, constrained binding (*3*, *38*, *49*, *50*). We suggest that LIMD1 scaffolding of AGO2–TNRC6 prolongs dwell time and helps miRISC overcome kinetic or structural barriers *in vivo*, potentially facilitating lateral scanning and supplementary pairing (*3*, *50–52*). In this framework, consistent with models in which weaker seeds rely more on supplementary pairing (*53–56*), GC-poor seeds show greater LIMD1-dependence; their intrinsic instability may permit AGO2 lateral scanning, with LIMD1 adapting AGO2 to extend dwell time and enable supplementary contacts (*3*, *50*). By contrast, GC-rich “stronger” seeds are sufficiently stable to bind with lesser LIMD1 dependence and often without supplementary contacts.

At a structural level, LIMD1 clamps AGO2 and TNRC6A through distinct domains (*19*). Whether this enhances AGO2–miRNA:target affinity by inducing specific conformations, reducing dissociation, or stabilising multivalent assemblies remains unresolved. The increased number of AGO2-bound miRNAs in LIMD1-deficient cells may reflect compensatory loading or reduced selectivity. These testable possibilities, together with limitations including incomplete paralog coverage (AGO/TNRC6), reliance on immortalised hSAECs, chimeric-eCLIP-intrinsic biases, and lack of direct biochemical, structural, or translational validation, warrant future mechanistic studies. Nonetheless, LIMD1 deficiency clearly compromises AGO2–miRNA targeting and repression, establishing AGO adaptors as essential architects of functional miRNA networks.

The implications extend beyond cell culture. LIMD1 LOH is frequently clonal in NSCLC and associates with reduced expression and poor survival (*28*, *29*, *33*). In lung tissue, LIMD1 inversely correlates with a miRNA:target set elevated in tumours compared to matched NAT, and this signature predicts LUAD survival. Together with *in vivo* evidence linking miRNA regulation to alveolar stem cell fate and oncogenic restraint (*57*), these findings position LIMD1 as a central determinant of lung homeostasis and tumour suppression. LIMD1 thus emerges as a guardian of miRNA network integrity, with LIMD1 haploinsufficiency representing a potential early prognostic marker in lung cancer.

Our evolutionary analyses reveal that younger, mammalian-specific miRNAs and less conserved target-sites are disproportionately LIMD1-dependent; these sites frequently showed accelerated evolution, formed stable duplexes, and concentrated within C_2_H_2_-ZNF transcripts, predominantly in CDS but also within 3′UTRs (*26*, *44*, *45*). Although the timing of LIMD1’s co-option into miRISC remains unknown, its integration appears to have broadened AGO2–miRNA targeting beyond conserved interactions to incorporate newly evolved miRNAs and sites, reshaping regulatory networks during metazoan evolution. Greater LIMD1 dependence at less conserved sites within individual miRNA repertoires supports this, showing that with LIMD1 conserved miRNAs can engage newly evolved targets and broaden their regulatory scope. The strong LIMD1 dependence of accelerated sites within C_2_H_2_-ZNFs, expanded in tetrapods and extensively diversified in mammals (especially primates and humans) (*44*, *45*, *58*, *59*), suggests that LIMD1 enabled AGO2– miRNAs to regulate new sites in this rapidly expanded TF family, linking adaptor-mediated miRISC function to the evolutionary expansion of regulatory complexity. LIMD1’s integration into miRISC thus represents a pivotal molecular innovation, exemplifying how adaptor proteins have shaped the diversification of post-transcriptional regulation in higher eukaryotes.

## Materials and Methods

### Cell culture

Primary-derived hSAECs (Lonza, CC2547) were maintained in hSAEC basal growth medium with supplements (Sigma-Aldrich, C-21170) at 37 °C in 5% CO₂ without antibiotics. Medium was refreshed every 2–3 days. At ∼70–80% confluence, cells were washed in PBS, detached with Trypsin/EDTA (Gibco, R001100) for ∼3 min, neutralized with DMEM (Sigma, D8437) containing 10% FBS (Gibco, 10270106), pelleted at 300 g for 3 min, and resuspended in fresh hSAEC medium. Cell counts were determined using a Countess II Automated Cell Counter (Invitrogen), and all cultures were routinely screened for mycoplasma. WT hSAECs were semi-immortalized by pFLRu-Bmi-1 transduction and subsequent puromycin selection.

### CRISPR-Cas9 LIMD1 editing and validation

To create LIMD1 Het and KO hSAECs, WT hSAECs were edited using the Dharmacon Edit-R CRISPR–Cas9 platform (Horizon Discovery), comprising EGFP–Cas9 mRNA (CAS11860), Edit-R tracrRNA (U-002005) and a predesigned crRNA targeting exon 1 of LIMD1 (guide #3; 5′-CCGAGTTTGAGGAAACTCGC-3′). A non-targeting synthetic sgRNA (#2; 5′-*propritary sequence*-3′; U-007502-01-05) served as negative control (to create Ctrl hSAECs). Transfections were performed with DharmaFECT Duo (Horizon, T-2010) according to the manufacturer’s protocol. Twenty-four hours after transfection, cells were seeded at limiting dilution (one cell per well) into 96-well plates for single-cell clonal expansion. Colonies were expanded after two weeks and transferred to replicate plates, where individual clones were screened by immunoblotting to identify potential heterozygous (Het) and knockout (KO) genotypes alongside controls (Ctrl).

LIMD1 mutations in candidate Het and KO clones were verified by PCR amplification of the targeted exon 1 locus, followed by Sanger sequencing (SourceBioscience). PCR products were generated with Phusion HF master mix (NEB, M0531S) and primers 5′-GAGTAGAGGCCCTGTCAATGG-3′ (forward) and 5′-CACAGATCCCAGGCTACCATC-3′ (reverse), purified, and sequenced using universal T7/SP6 primers.

WGS was subsequently performed to confirm on-target editing and to assess potential off-target effects that could impact miRNA-mediated regulation. Genomic DNA was extracted, quantified (NanoDrop OD260/280 = 1.8–2.0), and libraries were prepared by Novogene with ∼350 bp inserts, Illumina adapters, and limited-cycle PCR enrichment. Sequencing was performed on an Illumina NovaSeq 6000 with 150 bp paired-end reads, targeting ∼30× mean coverage across the GRCh38 reference genome. Reads were filtered for quality, aligned with BWA-MEM, duplicates marked with Picard, and variants called using GATK HaplotypeCaller. Putative off-target events were identified by local alignment of all detected variants against the 23 nt guide sequence plus PAM, requiring ≥13 nt contiguous seed match, ≤3 mismatches, and ≥10 supporting reads. No high-confidence off-target edits were detected.

### Western blotting

Cells were lysed in RIPA buffer (150 mM NaCl, 1% (v/v) IGEPAL, 0.5% (w/v) deoxycholic acid, 0.1% (w/v) SDS, 50 mM Tris-HCl, pH 7.5) supplemented with protease (Pierce, A32963) and phosphatase inhibitors (Roche, 12352204) during the log phase of growth. Protein concentration was quantified using the Pierce BCA Protein Assay Kit (Thermo Fisher Scientific, 23225). Equal amounts of protein lysate (15–30 µg per lane) were resolved by SDS–polyacrylamide gel electrophoresis (SDS–PAGE), transferred onto PVDF membranes, and probed with the following primary antibodies: mouse anti-LIMD1 (in-house, clone 3F2 C6); mouse anti-β-actin (Sigma, A1978); mouse anti-Vinculin (Invitrogen, 14-9777-82); rabbit anti-AGO2 (Sino Biological, 50683-RP02); rabbit anti-GW182/TNRC6A (Bethyl, A302-329A); rabbit anti-TNRC6B (Sigma, AB9913). HRP-conjugated secondary antibodies (Dako, P0214) were used in combination with enhanced chemiluminescence reagents (Thermo Fisher Scientific; Millipore, WBKLS0500). Blots were imaged using a GE Healthcare ImageQuant system. Densitometry of immunoblot band intensity was performed in ImageJ, with target signal intensities normalized to the Vinculin loading control.

### Co-immunoprecipitation (Co-IP)

Cells from a 15 cm dish (one dish per elution) were lysed in 1 mL NP-40 buffer (150 mM NaCl, 50 mM Tris-HCl, pH 8.0, 0.7% NP-40, 5% glycerol) supplemented with protease and phosphatase inhibitors. Lysates were clarified by centrifugation at 14,000 × g for 5 min at 4 °C. One millilitre (∼0.5 mg total protein) of cleared lysate was incubated with 1 µg of antibody (mouse IgA isotype control, eBioscience/Invitrogen, 14-4762-81; or mouse IgA anti-AGO2, Santa Cruz, sc-53521) overnight at 4 °C with rotation. For inputs, 80 µL of cleared lysate was mixed with 20 µL of 5× Laemmli buffer and boiled for 5 min. Antibody– lysate mixtures were washed twice in 500 µL NP-40 buffer and resuspended in 30 µL NP-40 buffer. Protein L magnetic beads (Pierce, 88849) were then added and incubated for 1 h at room temperature with rotation to capture antibody–protein complexes. Beads were washed four times in 1 mL PBS containing 0.1% Tween-20. Protein complexes were eluted by addition of 40 µL 0.2 M glycine (pH 2.5) with 3 min incubation at room temperature, with regular mixing. Eluates were transferred to new tubes and neutralized with 5 µL 1 M Tris-HCl (pH 8.0). Following addition of 11.25 µL 5× Laemmli buffer and boiling for 5 min, samples were analyzed by western blotting as described above. Densitometry was performed in ImageJ, with co-IP band intensities normalized to the relative input control (AGO2).

### Proximity ligation assay (PLA) and analysis

hSAECs grown on chamber slides (Nunc™ Lab-Tek™ II CC2™ Chamber Slide System; Thermo Fisher Scientific, 154852) were fixed with 4% paraformaldehyde in PBS for 15 min at room temperature, washed, and permeabilized according to the manufacturer’s instructions. PLA was then performed following the Duolink® PLA Fluorescence Protocol (Sigma-Aldrich) using mouse anti-AGO2 (Merck Millipore, 04-642; 1:400) and rabbit anti-GW182/TNRC6A (Bethyl, A302-329A; 1:300) as primary antibodies.

For negative controls, each primary antibody was paired with a species-matched IgG isotype control (rabbit mAb IgG, CST #3900; mouse mAb IgG2a, CST #61656), or a probes-only condition was used. After counterstaining with Phalloidin-AF488 (Cell Signaling, #8878) to define cell boundaries and DAPI to mark nuclei, coverslips were mounted in Duolink® Mounting Media.

Slides were imaged using a ZEISS LSM 880 confocal microscope with Airyscan (64× oil objective). Signal gain and offset were optimized for each experiment relative to negative controls. Z-stack images were processed using ZEN 3.4 (blue edition, Zeiss), ImageJ, and CellProfiler, with the latter used for per-cell quantification of uniformly thresholded PLA puncta (cell boundaries defined from F-actin outline and nuclei from DAPI). PLA interaction signal was quantified in ≥8 cells per condition across multiple fields and wells (≥3), and images were digitally adjusted for clarity in figures without altering relative intensities.

### Psi-Check-2 cloning and Dual-Luciferase reporter assays

Oligonucleotides encoding empty vector (VO), tandem miRNA sites with seed matches (T) or mismatches (NT), or artificial reporters were cloned into the 3′ UTR region of the Renilla luciferase cassette in psiCheck-2 (Promega, C8021) using XhoI and NotI restriction sites. Tandem miR-99/100 sites (T: 5′-AGCAAGTGTAACGG**TACGGGT**A-3′; NT: 5′-AGCAAGTGTAACGG**TA***ATAAC*A-3′) contained five repeats and were created as previously described (*19*). Artificial reporters included a let-7a construct containing six seed-matched sites (5′-AACTATACAACGT**CTACCTC**A-3′) and a miR-21 construct containing three seed-matched sites (5′-TCAGAAG**ATAAGCT**AGGGGTCA-3′; 5′-TCAGAAG**ATAAGCT**AGGGGTCA-3′; 5′-TCAGAAG**ATAAGCT**AGTTTAAA-3′). Bold indicates the seed-matched region. Constructs were verified by colony PCR, diagnostic XhoI/NotI digest, and Sanger sequencing, and plasmids were diluted to 100 ng/µL in TE buffer for transfection.

For luciferase assays, 5,000 cells per well were seeded in 96-well plates and transfected 24 h later with 20 ng psiCheck-2 plasmid using Attractene (QIAGEN, 301005) in Opti-MEM (Thermo Fisher Scientific, 31985070). After 24 h, cells were lysed in 25 µL Passive Lysis Buffer (Promega, E1960) and subjected to one freeze–thaw cycle. Lysates (20 µL) were transferred to white-walled plates for sequential luminescence measurement on a FLUOstar Omega plate reader (BMG Labtech) with 25 µL Firefly substrate followed by 25 µL Renilla substrate. Each condition was assayed in ≥4 technical replicates across ≥3 independent experiments. Renilla activity was normalized to Firefly internal control and expressed relative to VO to calculate miRNA-mediated repression.

### Chimeric miR-eCLIP protocol

Chimeric miR-eCLIP was performed as described previously (*32*). Briefly, 2 x 1.25 million cells from four samples per group (three independent CRISPR clones and one clone repeated) were seeded into 15 cm dishes and grown to ∼80% confluence (∼2 x 10 million cells each) before UV-crosslinking on ice at 254 nm (400 mJ/cm²). Cells were scraped, pelleted at 300 g, flash-frozen in liquid nitrogen, and stored at –80 °C. Pellets were lysed in cold eCLIP buffer containing protease and RNase inhibitors, then sonicated on a QSonica Q800R (75% amplitude, 30 s on/off for 10 cycles). Lysates were adjusted to fragment RNA with RNase I according to sample RIN, clarified by centrifugation, and aliquoted to yield 20 µg RNA per IP. Immunoprecipitation was carried out overnight at 4 °C with AGO2 antibody (Santa-Cruz, sc-53521)–coated magnetic beads (M280 Sheep Anti-Mouse IgG Dynabeads, Thermo Fisher 11202D). Beads were washed under stringent high-salt and no-salt conditions, sequentially treated with PSP and PNK enzymes to repair RNA ends, and subjected to on-bead chimeric miRNA:mRNA ligation with T4 RNA ligase. Following additional washes, 3′ RNA adapters were ligated on-bead and RNA was released by proteinase K digestion.

Matched Input RNA samples were processed in parallel: column cleanup, PSP/PNK end repair, and 3′ adapter ligation. Both IP and Input RNAs were reverse-transcribed using a chimera-specific primer, and the resulting cDNA underwent end repair by nuclease treatment and mild alkaline hydrolysis with bead-based cleanup. Overnight ligation of a single-stranded cDNA adapter was followed by PCR amplification using Illumina i7/i5 index primers, with cycle numbers determined by qPCR. Final libraries were purified with AMPure XP beads, quality-checked on an Agilent TapeStation, pooled equimolarly, and sequenced single-end (100 bp) on an Illumina platform, targeting ∼50 million reads per IP sample and ∼40 million reads per Input sample. Further details on chimeric miR-eCLIP protocol can be obtained upon request.

### Chimeric miR-eCLIP analysis

Initial processing and analysis was performed by Eclipsebio with a proprietary analysis pipeline v1 developed from several published eCLIP publications (*32*, *60*, *61*). The original bioinformatic pipelines are available (https://github.com/yeolab/eclip; https://github.com/YeoLab/chim-eCLIP). Briefly, UMIs were pruned with umi_tools (v1.1.1), and 3′ adapters were trimmed with cutadapt (v3.2). Reads <18 nt after trimming were discarded. Reads were first aligned to a database of repetitive elements and rRNA sequences; non-repeat reads were then mapped to the human genome (GRCh38/hg38, UCSC) using STAR (v2.7.7a). PCR duplicates were removed with umi_tools. AGO2 eCLIP Read clusters were identified with CLIPper (v2.0.1) (*62*), and IP versus Input fold enrichment and statistical significance were calculated; clusters meeting predefined thresholds log₂ fold enrichment ≥ 3 and p ≤ 0.001 were designated as AGO2–peaks.

In parallel, non-chimeric reads were used to quantify AGO2-bound miRNA abundance. Reads were adapter-trimmed, UMI-deduplicated, and aligned to mature miRNA sequences from miRBase (v22.1) using Bowtie, allowing multimapping to account for homologous family members. Abundance was expressed as reads per million (RPM) relative to the total AGO2-bound miRNA read count per sample. These data enabled direct comparison of miRNA expression levels between Ctrl, Het, and KO conditions.

Unmapped reads were “reverse mapped” to mature miRNAs from miRBase (v22.1) using bowtie (v1.2.3). The miRNA sequence was trimmed, and the remainder was aligned to the genome with STAR; PCR duplicates were again removed with umi_tools. miRNA:target clusters were identified with CLIPper and annotated with the responsible miRNA(s). Peaks were annotated using transcript information from GENCODE release 41 (GRCh38.p13) with priority given to protein-coding features (CDS, UTRs, introns) over non-coding transcripts (exons, introns). AGO2 clusters were normalized to paired Input samples and reproducibility was assessed by IDR (Irreproducible Discovery Rate) analysis. Reproducible chimeric AGO2–miRNA:target peaks were defined as those with ≥3 chimeric reads from the same miRNA seed family overlapping the peak in each replicate.

Each reproducible chimeric peak was therefore treated as a unique AGO2–miRNA:target-site interaction with its seed match. After UMI de-duplication, each retained chimeric read reflects a unique ligation event captured in the experiment, providing a measure of relative interaction abundance rather than absolute molecular counts (*32*). While the low sensitivity of ligation assays means many true interactions are not recovered, and ligation or sequencing biases may influence capture, UMI de-duplication ensures counts are not inflated by PCR artifacts.

Peak-level data (genomic coordinates, gene, RNA feature, enrichment/chimeric read counts, fold changes, and p-values) were further analyzed using Microsoft Excel and GraphPad Prism v9.5.1. Additional details of the analysis pipeline are available upon request. AGO2-miR-eCLIP data (input and IP) file (.bigWig) have been deposited to GEO under the accession number GSE304955.

### Positionally enriched k-mer analysis (PEKA) analysesh

PEKA was applied as described previously (*36*) to reproducible AGO2–miRNA chimeric peaks. Putative crosslink sites (Xn) were partitioned into thresholded sites (tXn; within peaks containing ≥70% of the regional cDNA signal) and reference sites (oXn; outside peaks), with all others discarded. For each set, sequences flanking tXn and oXn (±100 nt) were extracted, scanned for k-mer presence, and used to calculate relative occurrence (RtXn). RtXn scores represent fold-enrichment of a k-mer at each position relative to flanking background (RtXn = 0: no binding; 1: background level; >1: position-specific enrichment). The PEKA score was defined as the standardized difference between a k-mer’s mean RtXn at enriched positions and its mean RtXn from randomly sampled background sites, capturing both the magnitude and positional consistency of enrichment.

For visualization and quantitative comparison across Ctrl, Het, and KO conditions, clustered heatmaps of the top 40 k-mers by PEKA score were generated. From these data, additional metrics were derived, including: (i) GC content of the top 40 motifs; (ii) maximal occurrence values, defined as the highest enrichment at centered, upstream, or downstream positions relative to the crosslink site, as indicated by dot positioning on the PEKA heatmap; (iii) mean enrichment per nucleotide position, calculated by averaging enrichment across motifs at each position relative to tXn within each condition; Gaussian smoothing (σ = 3.33) was applied to reduce noise and generate positional enrichment profiles; (iv) local Shannon entropy, computed in sliding 9×9-pixel windows across motif heatmaps, with mean entropy values calculated per motif row. Entropy quantifies variability in positional enrichment, with higher values reflecting smoother, more distributed footprints and lower values reflecting punctate binding profiles. Together, PEKA analysis provides positional resolution of target-motif sequences relative to the AGO2 crosslink site, enabling assessment of how LIMD1 deficiency alters motif positioning around AGO2. All code (PEKA, entropy, kernel density plotting) is available upon request.

### Canonical seed GC content calculation and matched canonical peak analysis

Canonical seed matches (8mer, 7mer-m8, 7mer-A1, 6mer) were identified from chimeric peaks. For each, the corresponding miRNA sequence was retrieved from miRBase (mature.fa), and the 6-nt seed (positions 2–7) was extracted. GC content was calculated as the proportion of G/C bases and used to classify interactions as GC-poor, intermediate, or GC-rich. Matched canonical interactions were here defined as those detected across all conditions with overlapping genomic coordinates (±10 bp). For each matched peak, mean chimeric read counts were obtained per condition, and percentage changes relative to Ctrl were calculated after stratification by GC category.

### mRNA-seq library preparation and analysis

hSAECs (1.25 million cells per 15-cm dish; one dish per CRISPR clone, three clones per group) were cultured for 7 days (media changed three times) to ∼80% confluence (∼10 million cells/dish). After pelleting, total RNA was extracted from frozen pellets using the miRNeasy Mini Kit (Qiagen), and poly(A) RNA was enriched with Dynabeads Oligo(dT)25 (Thermo Fisher). RNA integrity was assessed on an Agilent 4200 TapeStation.

For mRNA-seq libraries, 50 ng poly(A) RNA was fragmented in 2× FastAP buffer (Thermo Fisher), dephosphorylated with FastAP, phosphorylated with T4 PNK (NEB), and ligated at the 3′ end to an RNA adapter with T4 RNA ligase (NEB). After Zymo column cleanup, fragments were reverse-transcribed with SuperScript III (Invitrogen), treated with ExoSAP-IT (Affymetrix), and ligated at the cDNA 3′ end to a 5′ Illumina adapter. Libraries were cleaned on Dynabeads MyOne Silane (Thermo Fisher). Cycle number was determined by qPCR, followed by final amplification with barcoded primers (Q5 polymerase, NEB) and size selection using AMPure XP beads (Beckman). Libraries were quantified on the TapeStation and sequenced on an Illumina NovaSeq.

Reads containing a 10-nt UMI at the 5′ end were processed with umi_tools (v1.0.1), adapter-trimmed with cutadapt (v2.7), aligned first to RepBase 18.05 then to hg38 using STAR (v2.7.6a), and deduplicated by UMI and position with umi_tools. Gene counts were generated against GENCODE v29 annotations, and differential expression was analyzed using DESeq2 (v1.34.0). Bulk mRNA-seq data (csv files containing unnormalised raw gene counts) have been deposited to GEO under the accession number GSE305016.

### Total Proteomics

Proteomics experiments were performed using mass spectrometry (MS) as previously reported (*63*, *64*), with minor technical modifications, on six independently plated, lysed, and processed replicates of Ctrl C4, Het C6, and LIMD1 KO C2 hSAECs. Briefly, cells were seeded and grown out in 15 cm dishes for 7 days and pelleted as described as described for chimeric eCLIP. Frozen pellets were lysed in 8 M urea buffer (Sigma-Aldrich) and supplemented with phosphatase inhibitors (10 mM Na_3_VO_4_, 100 mM β-glycerol phosphate, 25 mM Na_2_H_2_P_2_O_7_ ((Sigma-Aldrich)). Proteins were digested into peptides using trypsin as previously described (*65*, *66*). From each sample, 30 μg of protein was reduced and alkylated sequentially with 10 mM dithiothreitol (1 h, 25 °C, agitation) and 16.6 mM iodoacetamide (30 min, 25 °C, dark). Trypsin beads (50% slurry of TLCK-trypsin; ThermoFisher) were equilibrated in 20 mM HEPES (pH 8.0), the urea concentration was reduced to 2 M, and 70 μL beads were added for overnight digestion at 37 °C. Beads were removed by centrifugation (2000 g, 5 min, 5 °C). Peptides were desalted using the AssayMAP Bravo platform (Agilent Technologies, peptide clean-up v3.0). Reverse phase S cartridges (Agilent, 5 μL bed volume) were primed with 250 μL 99.9% acetonitrile (ACN) with 0.1% TFA and equilibrated with 250 μL 0.1% TFA at a flow rate of 10 μL/min. The samples were loaded at 20 μL/min, followed by an internal cartridge wash with 0.1% TFA at a flow rate of 10 μL/min. Peptides were then eluted with 105 μL of solution (70/30 ACN/H2O + 0.1% TFA). Eluted peptide solutions were dried in a SpeedVac vacuum concentrator and stored at −80 °C.

Samples were run in the LC-MS system over consecutive days to reduce technical variability. Peptide pellets were reconstituted in 5 µL of 0.1% TFA, after which 2 µL of this solution were further diluted in 18 µL of 0.1% TFA: 2 µL were injected into the LC-MS/MS system. The LC-MS/MS platform consisted of a Dionex Ultimate 3000 RSLC coupled to Q Exactive™ Plus Orbitrap Mass Spectrometer (Thermo Fisher Scientific) through an EASY-Spray source. Mobile phases for the chromatographic separation of the peptides consisted of Solvent A (3% ACN; 0.1% FA) and Solvent B (99.9% ACN; 0.1% FA). Peptides were loaded in a μ-pre-column and separated in an analytical column using a gradient running from 3 to 28% B over 90 min. The UPLC system delivered a flow of 10 µL/min (loading) and 250 nL/min (gradient elution). The Q Exactive Plus operated a duty cycle of 2.1 s. Thus, it acquired full scan survey spectra (m/z 375–1500) with a 70,000 Full Width at Half Maximum (FWHM) resolution followed by data-dependent acquisition in which the 15 most intense ions were selected for HCD (higher energy collisional dissociation) and MS/MS scanning (200–2000 m/z) with a resolution of 17,500 FWHM. A dynamic exclusion period of 30 s was enabled with m/z window of ±10 ppm. MS data collection was carried out using Thermo Scientific FreeStyle 1.4.

MS raw files were converted into Mascot Generic Format using Mascot Distiller (version 2.8.5) and searched against the UniProt database (Uni Prot.fasta) restricted to human entries using the Mascot search daemon (version 3.0.0). Allowed mass windows were 10 ppm and 25 mmu for parent and fragment mass to charge values, respectively. Identified peptides were quantified using in-house software Pescal as described previously (*65*). Quantitative data were processed in R (v4.2.2), Excel, and GraphPad Prism v9.5.1. Only peptides identified in all replicates of all groups and mapping uniquely to proteins were included in final analyses (n = 2667). The analysis code is available at https://github.com/CutillasLab/protools2.

### Gene-set selection and lung expression and survival analysis for LIMD1–AGO2– miRNA:target signature analyses

To define a LIMD1–AGO2:miRNA-target signature we hypothesised to be dysregulated in LIMD1-deficient lung adenocarcinoma, we applied the following criteria: in both Het and KO, genes with significantly de-enriched 3′UTR chimeric peaks or within significantly dysregulated targetomes of five miRNAs; increased mRNA expression (q < 0.05); increased protein levels (p < 0.1); and LIMD1 dose-dependency of chimeric read loss, mRNA increase, and protein increase. Five genes met these criteria: *VDAC1*, *HMGB2*, *KIF2A*, *IGF2BP3*, and *PPIF*. To avoid bias, (i.e., only then further analysing genes which showed desired results), no other or further selections were performed.

Using TCGA LUAD mRNA data (normalized log₂[FPKM-UQ + 1]; n = 517 tumour, n = 72 normal samples) and CPTAC mRNA and protein datasets (n = 99 matched LUAD tumour– NAT paired samples) (*67*), Z-scores for the five-gene set were computed with the R package GSVA; NAT and tumour expression values were used as the reference. These Z-scores were then correlated with LIMD1 mRNA or protein abundance to generate signature Pearson correlations. Survival analyses (Kaplan–Meier, univariate, and multivariate Cox proportional hazards) were performed using TCGA LUAD overall survival and clinical annotation data. The composite score (TCGA LUAD) was calculated as z-transformed LIMD1 mRNA expression minus the z-transformed target set expression. Patients were dichotomized by upper and lower quartiles (highest vs lowest) for Kaplan–Meier survival analyses. Protein-level survival analysis was not performed due to the absence of public MS-protein survival datasets.

### Evolutionary conservation of miRNAs, target-sites and thermodynamic stability of miRNA:target duplexes

The ages of miRNA loci and families were obtained from miRGeneDB 2.1 (*26*) and species divergence dates were taken as the median values reported in TimeTree 5 (*68*). LIMD1-dependent canonical interactions were defined as those showing reduced enrichment (p < 0.3) in chimeric eCLIP data in KO versus Ctrl. Logistic regression models were constructed as: [retained/lost in LIMD1 KO] ∼ expression_level + conservation + interaction. Expression level was defined from average miRNA abundance in Ctrl cells and classified into three groups: highly expressed (top 10%), lowly expressed (bottom 10%), and mid-level (remaining 80%). Mature miRNA sequences were obtained from miRBase v22 (*69*). Canonical target predictions were generated with seedVicious v1.1 (*43*), which was also used to compute predicted miRNA:target duplex hybridization energies using the ViennaRNA package (*70*). Conservation of AGO2–miRNA:target sites was measured as PhyloP scores (*42*) at the first position (3′) of the target seed match (sixmer), derived from the vertebrate 100-way alignment (hg38; https://hgdownload.soe.ucsc.edu/goldenPath/hg38/phyloP100way/.). GC seed content of canonical interactions was calculated as described above.

### Gene ontology molecular function (GO: MF) analysis

Canonical LIMD1-dependent sites (reduced enrichment in KO vs. Ctrl; p < 0.3) with negative PhyloP scores were subjected to Gene Ontology molecular function (GO:MF) enrichment analysis using GOrilla (http://cbl-gorilla.cs.technion.ac.il), with all genes identified in mRNA-seq used as the background set. Enriched GO:MF terms were retained at false discovery rate (FDR) q < 0.001. To reduce redundancy, significant terms were clustered with REViGO (http://revigo.irb.hr). Final GO categories were grouped manually into DNA-binding/transcription-related and small-molecule/ion-binding functions. Genes encoding C_2_H_2_-ZNF proteins were assigned according to HUGO Gene Nomenclature Committee (HGNC) gene group annotations.

### Statistical analyses

Statistical analyses were performed after assessing data normality and selecting tests appropriate to experimental design and data structure. Multiple-comparison corrections were applied where required. Statistical values are reported in figures and legends. All analyses were conducted in GraphPad Prism v9.5.1 or Python, except for miR-eCLIP differential enrichment and mRNA-seq differential expression, which were analyzed with DESeq2 as described above.

## Acknowledgments

We thank all members of the Sharp laboratory and the Barts Cancer Institute core facilities for helpful discussions and technical support. We are also grateful to current and former members of the Cancer Cell and Molecular Biology Department at Barts Cancer Institute for sharing insight and reagents throughout, particularly Giulia Giuducci for discussions on experimental work and manuscript feedback; Emilie Alard, Martin Dodel, and Lovorka Stojic for discussions related to experimental work; and Natalie Bevan for feedback on the manuscript.

## Author Contributions

Conceptualization: A.F.F.C., K.M.S., D. L., S.G.J., A.M., T.V.S

Investigation: A.F.F.C., K.M.S., A.M., T.V.S.

Formal analysis: A.F.F.C., A.M., E.M., A.T., K.S., V.R., S.G.J., T.V.S.

Supervision: T.V.S., K.M.S.

Writing – original draft: A.F.F.C., T.V.S.

Writing – review & editing: all authors contributed to the writing, review and editing of the manuscript.

## AI-assisted writing and coding disclosure

Artificial-intelligence tools were used in limited aspects of manuscript preparation and computational work. OpenAI’s ChatGPT 5 Pro (version as of September 2025) was used to refine the clarity, tone, and structure of the manuscript text and figure captions, and OpenAI’s ChatGPT 5 Pro and Google’s Gemini were used to assist in code generation and troubleshooting for selected analyses. AI tools were not used to perform, interpret, or manipulate analyses, data, or figures, and no AI-generated images or multimedia were included. The following, or similar, prompt was provided during text editing: “Please help me refine the clarity, tone, and conciseness of the following scientific text without altering the meaning or scientific content. Maintain a formal academic tone appropriate for a high-impact journal.”

All scientific content, analyses, code, and conclusions were conceived, executed, and verified by the authors. The corresponding author takes full responsibility for the accuracy, originality, and integrity of the manuscript and confirms that all AI-assisted outputs were critically reviewed to ensure factual correctness and to eliminate any potential bias.

## Data and Materials Availability

All AGO2-miR-eCLIP and bulk mRNA-sequencing data have been deposited in GEO under accession number GSE304955 and GSE305016, respectively. These await journal publication before public access; access via token code can be requested. Additional data supporting the findings of this study (processed reproducible AGO2-miR-eCLIP peaks, mRNA-seq DESEq2 analyses, MS fold changes, canonical interaction seed GC-content, phyloP score, and predicted duplex stability) are available in the Supplementary Materials. Code for plotting analyses are available upon request.

