## Supplementary Figures and Legends for "An AGO2 adaptor expands the functional and evolutionary reach of microRNA targeting"

### Figure S1. LIMD1 potentiates AGO2–TNRC6A interactions and miRNA reporter repression

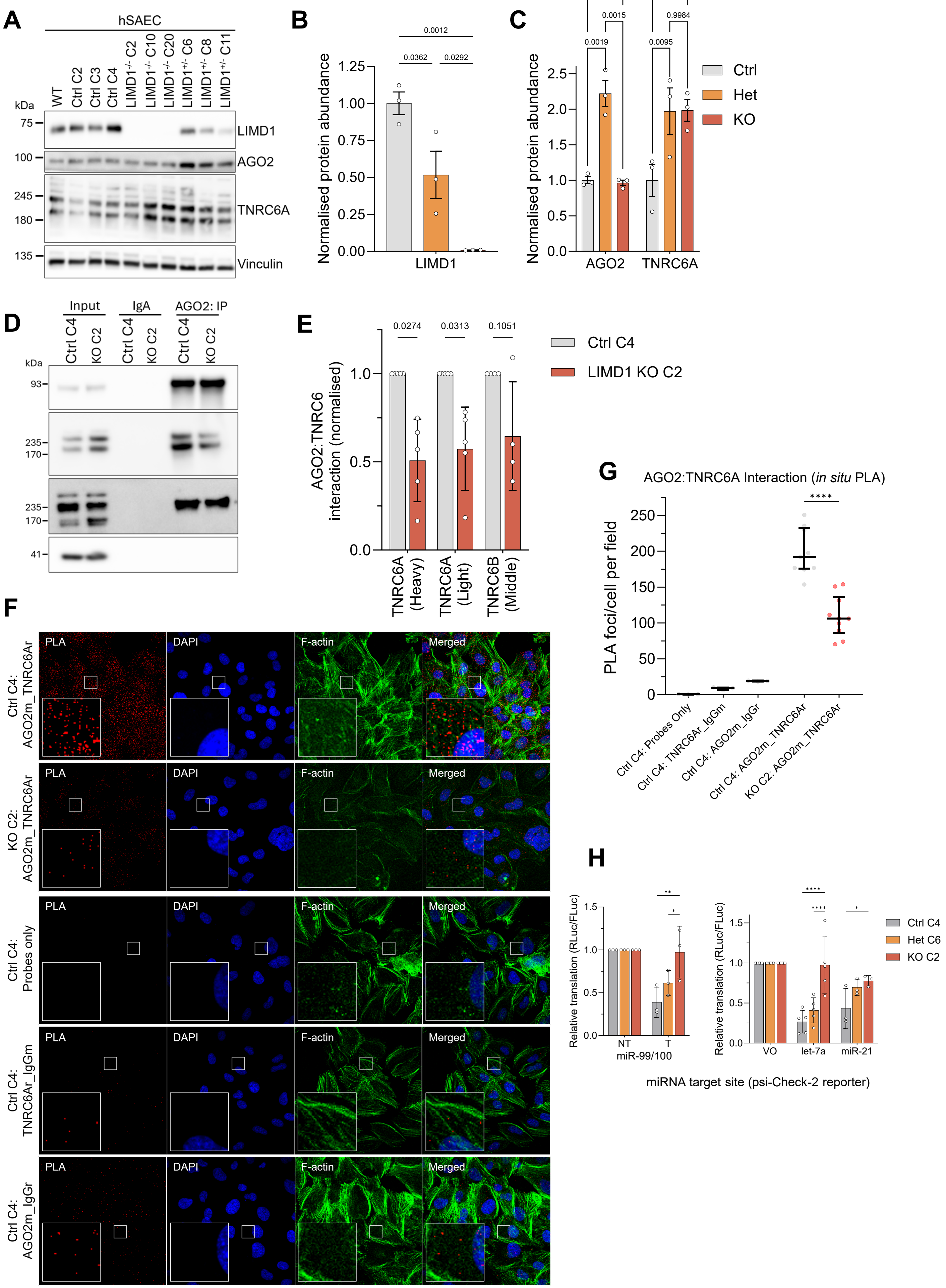

**Figure S1. LIMD1 potentiates AGO2–TNRC6A interactions and miRNA-reporter repression**

**(A)** CRISPR-Cas9 editing of hSAECs generated Ctrl ($LIMD1^{+/+}$), Het ($LIMD1^{+/-}$), and KO ($LIMD1^{-/-}$) clones (n=3 per genotype). **(B, C)** Immunoblot quantification of LIMD1 **(B)** and AGO2/TNRC6A **(C)** normalised to vinculin and Ctrl mean (mean ± SEM, n=3). LIMD1 was tested by one-way ANOVA; AGO2/TNRC6A by two-way ANOVA with multiple-comparisons correction. **(D, E)** Co-immunoprecipitation shows reduced AGO2–TNRC6A/B association in KO cells (mean ± SD, n≥4; unpaired t-test with correction). **(F, G)** *In situ* proximity ligation assay (PLA) reveals reduced AGO2:TNRC6A proximal association (< ~ 40nm) in LIMD1-deficient cells. **(F)** Representative fluorescence images showing PLA puncta (red), nuclei (DAPI, blue), and F-actin (green); negative controls included. Insets show 10× magnified regions of boxed areas. PLA channel thresholded for visual clarity. **(G)** Quantification of PLA foci per cell, averaged per imaging field (n≥3). Each dot represents one field. Statistical significance determined by Welch's two-tailed t-test on field means (*****P*<0.0001). **(H)** Dual-luciferase assays show reduced repression of psiCHECK-2 reporters bearing synthetic miRNA sites (miR-99/100, let-7a, miR-21) with a dose-dependent trend from Ctrl to Het to KO. Non-targeting (NT) or vector-only (VO) served as negative controls. Data are mean ± SD from ≥3 experiments; two-way ANOVA with correction; **P*<0.05, ***P*<0.01, *****P*<0.0001.

### Figure S2. LIMD1 deficiency increases AGO2–miRNA interactions

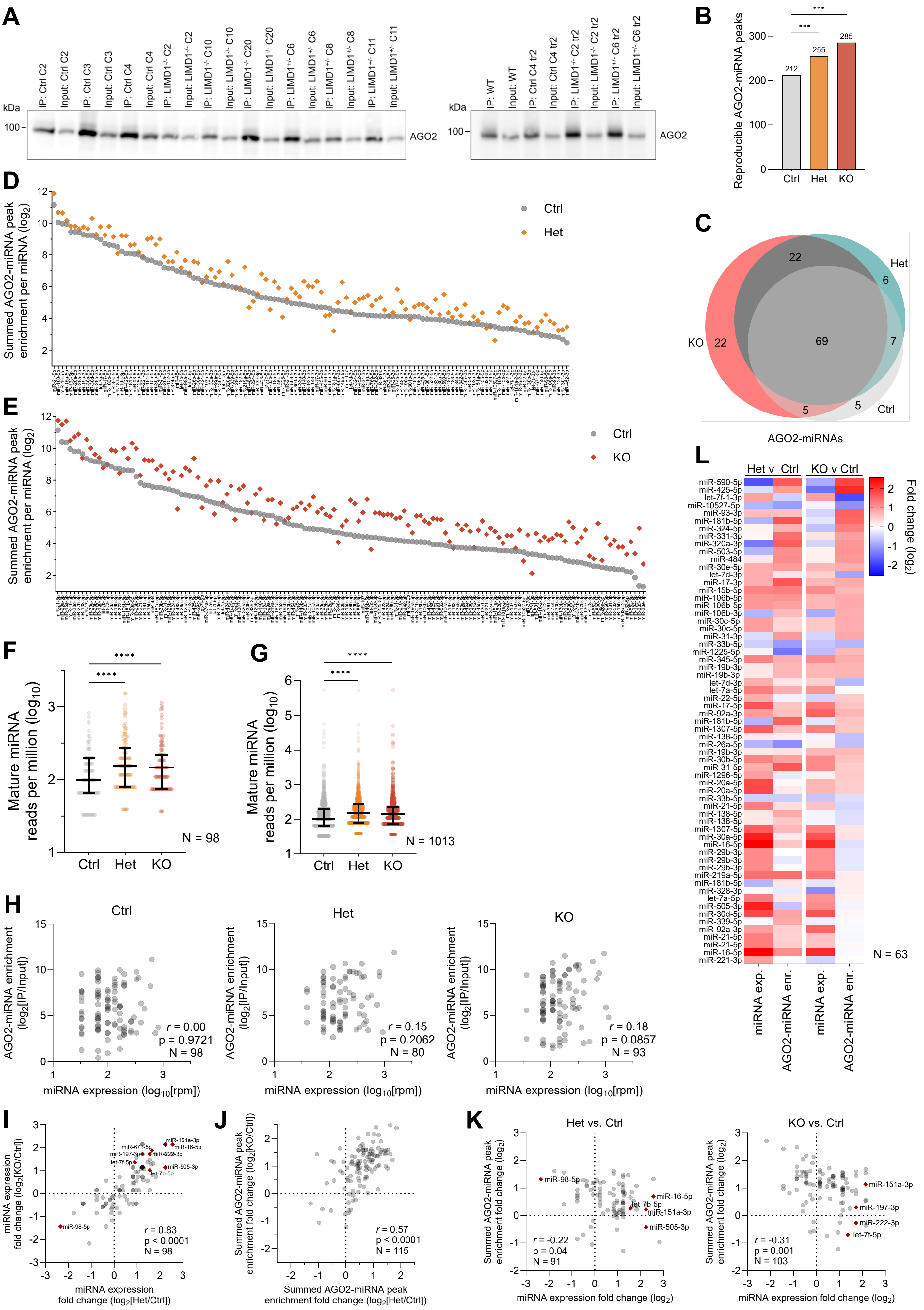

**Figure S2. LIMD1 deficiency increases AGO2–miRNA interactions independently of miRNA expression**

**(A)** AGO2-IP enrichment over input across chimeric-eCLIP replicates. **(B)** Dose-dependent increase in reproducible AGO2–miRNA peaks in LIMD1-deficient cells (pairwise proportion test, Benjamini–Hochberg correction; ***$P$<0.001). **(C)** Venn diagram of AGO2-associated miRNAs across Ctrl, Het, and KO. **(D, E)** Increased summed peak enrichment per AGO2–miRNA in Het and KO versus Ctrl. **(F, G)** Global increase in mature miRNA expression (reads per million, $\log_{10}$) for AGO2-associated miRNAs **(F)** and all mature miRNAs **(G)**. **(H)** Within groups, AGO2–miRNA enrichment correlates poorly with miRNA expression (Spearman), indicating regulation beyond expression levels. **(I, J)** Positive correlation of changes in miRNA expression (I) and summed AGO2–miRNA enrichment (J) in Het and KO versus Ctrl (Spearman), consistent with LIMD1-specific effects. **(K)** Moderate negative correlation between AGO2–miRNA enrichment and mature miRNA expression changes (Spearman), indicating expression alone does not drive enrichment. **(L)** Heatmap of $\log_2$ fold changes in mature miRNA expression and AGO2 peak enrichment, showing overall increases in both without clear relation to miRNA expression.

### Figure S3. AGO2–mRNA binding requires LIMD1

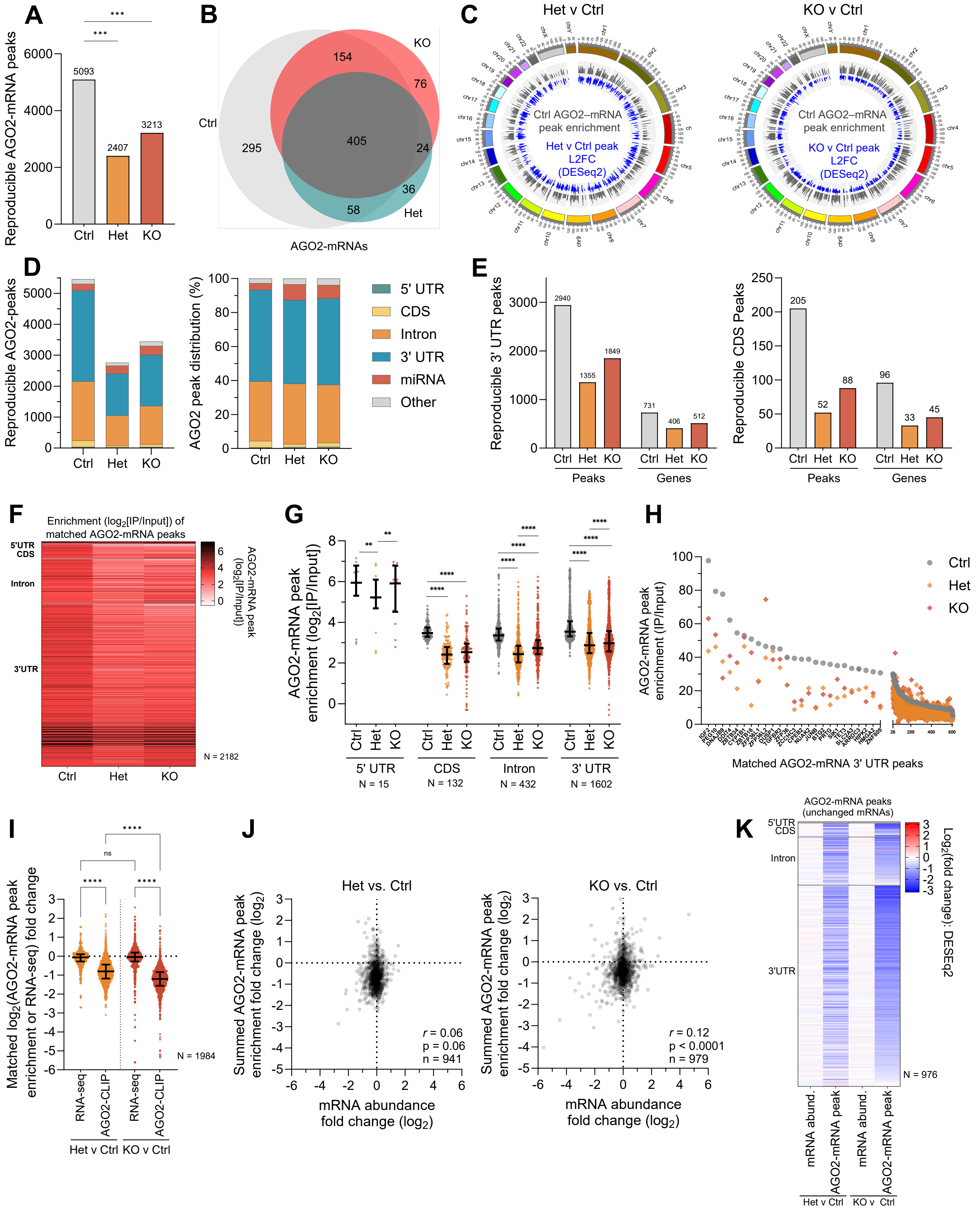

**Figure S3. AGO2–mRNA binding requires LIMD1**

**(A)** Fewer AGO2–mRNA peaks indicate reduced distinct AGO2–mRNA interactions in LIMD1-deficient cells (pairwise proportion test, Benjamini–Hochberg correction; **$P$<0.001). **(B)** Venn diagram of AGO2-associated mRNAs in Ctrl, Het, and KO, showing reduced repertoire in LIMD1-deficient cells. **(C)** Circos plot of global AGO2–mRNA peak enrichment: gray = Ctrl, blue = $\log_2$FC in Het or KO versus Ctrl. **(D)** Number and distribution of reproducible peaks by genomic feature (peaks hierarchically annotated as CDS, 5′UTR, 3′UTR, intron, miRNA, other). **(E)** Reproducible AGO2–mRNA peaks and distinct transcripts mapping to CDS's and 3′UTRs. **(F, G)** Enrichment ($\log_2$FC AGO2 IP/Input) for 2,182 matched AGO2–mRNA peaks, shown as a heatmap **(F)** and stratified by genomic region **(G)**. **(H)** Per-peak enrichment of 606 matched 3′UTR peaks; for transcripts with multiple peaks, only the most enriched is plotted. **(I)** AGO2–mRNA peaks show greater de-enrichment than transcript-level changes, indicating loss of binding is not driven by reduced mRNA abundance (Wilcoxon matched-pairs signed-rank test; ***$P$<0.0001). **(J)** Low correlation (Spearman) between mRNA abundance changes and summed AGO2–mRNA peak enrichment confirms major changes not driven by transcript levels. **(K)** Heatmap of $\log_2$ fold changes for mRNAs with stable abundance (0.8–1.2-fold, q>0.9), grouped by genomic feature and ordered by de-enrichment in KO, showing widespread peak loss among stable mRNA transcripts.

### Figure S4. LIMD1 deficiency constrains global AGO2–miRNA targeting

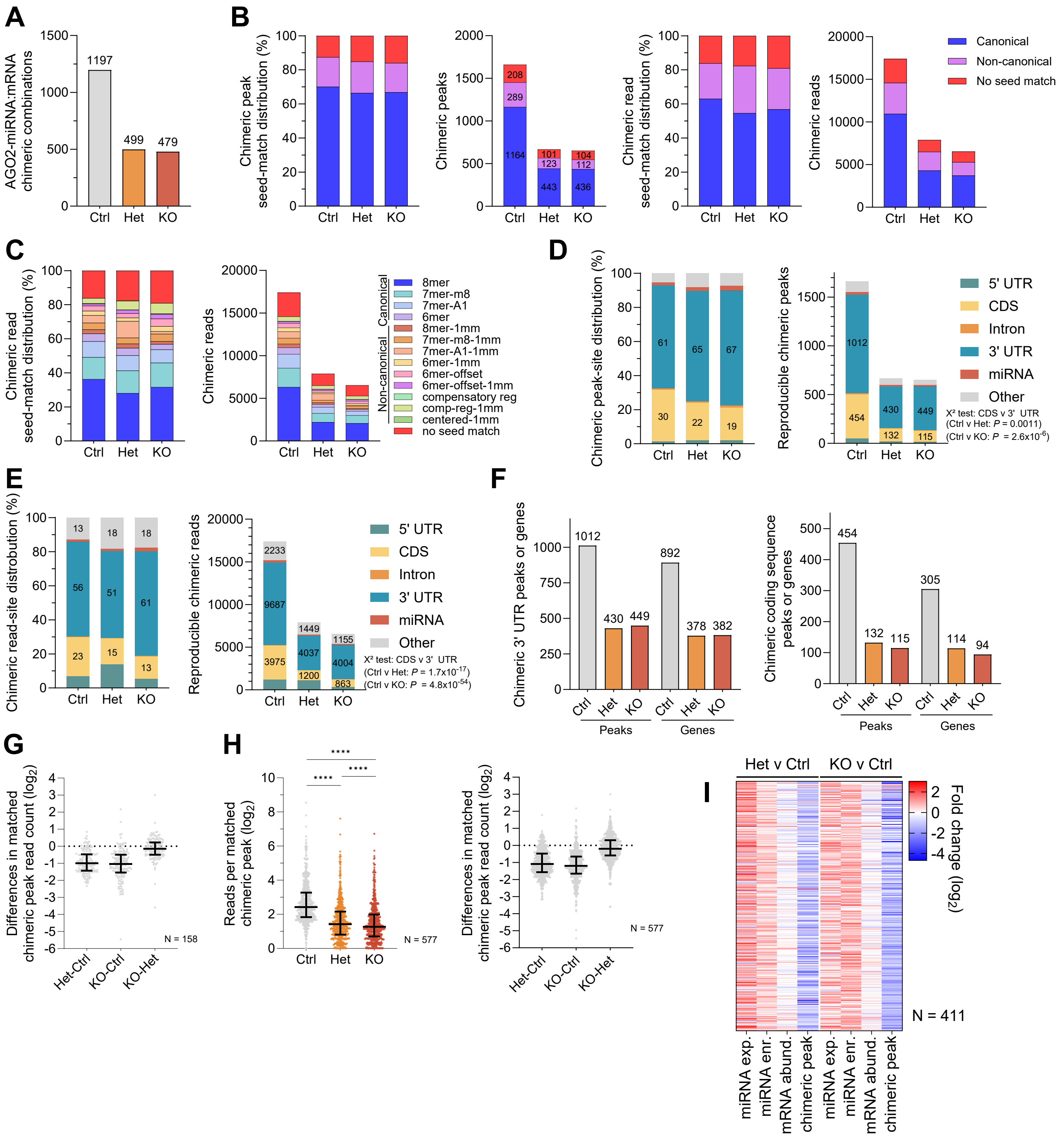

**Figure S4. LIMD1 deficiency constrains global AGO2–miRNA targeting**

**(A)** Fewer unique AGO2–miRNA:mRNA combinations in LIMD1-deficient cells. **(B)** Distribution and counts of chimeric peaks and reads by seed-match class (canonical, non-canonical, or no seed match; TargetScan definition). **(C)** Distribution of chimeric read counts by exact seed-match type (TargetScan definition) shows broad reductions across classes in LIMD1-deficient cells. **(D, E)** Genomic feature distribution of reproducible chimeric-target peaks **(D)** and reads **(E)**, annotated as CDS, 5′UTR, 3′UTR, intron, miRNA, or other ($\chi^2$ test for CDS vs 3′UTR peak distribution). **(F)** Fewer reproducible chimeric peaks and unique target transcripts mapping to CDS and 3′UTR target-sites. **(G)** Reduced normalized reads per matched chimeric peak (≥3 reads in Ctrl, Het, KO; Wilcoxon matched-pairs signed-rank test; ***$P$<0.0001). **(H)** Reduced normalized reads per matched chimeric peak (≥3 reads in any group, pseudo + 0.01 added to peaks with 0 reads to avoid $\log_0$ values; Wilcoxon matched-pairs signed-rank test; ***$P$<0.0001). **(I)** Heatmap of $\log_2$ fold changes in miRNA expression, AGO2–miRNA enrichment, mRNA abundance, and chimeric reads for matched chimeric peaks with complete data in Het and KO versus Ctrl.

### Figure S5. LIMD1 deficiency constrains breadth and depth of targeting by each AGO2–miRNA

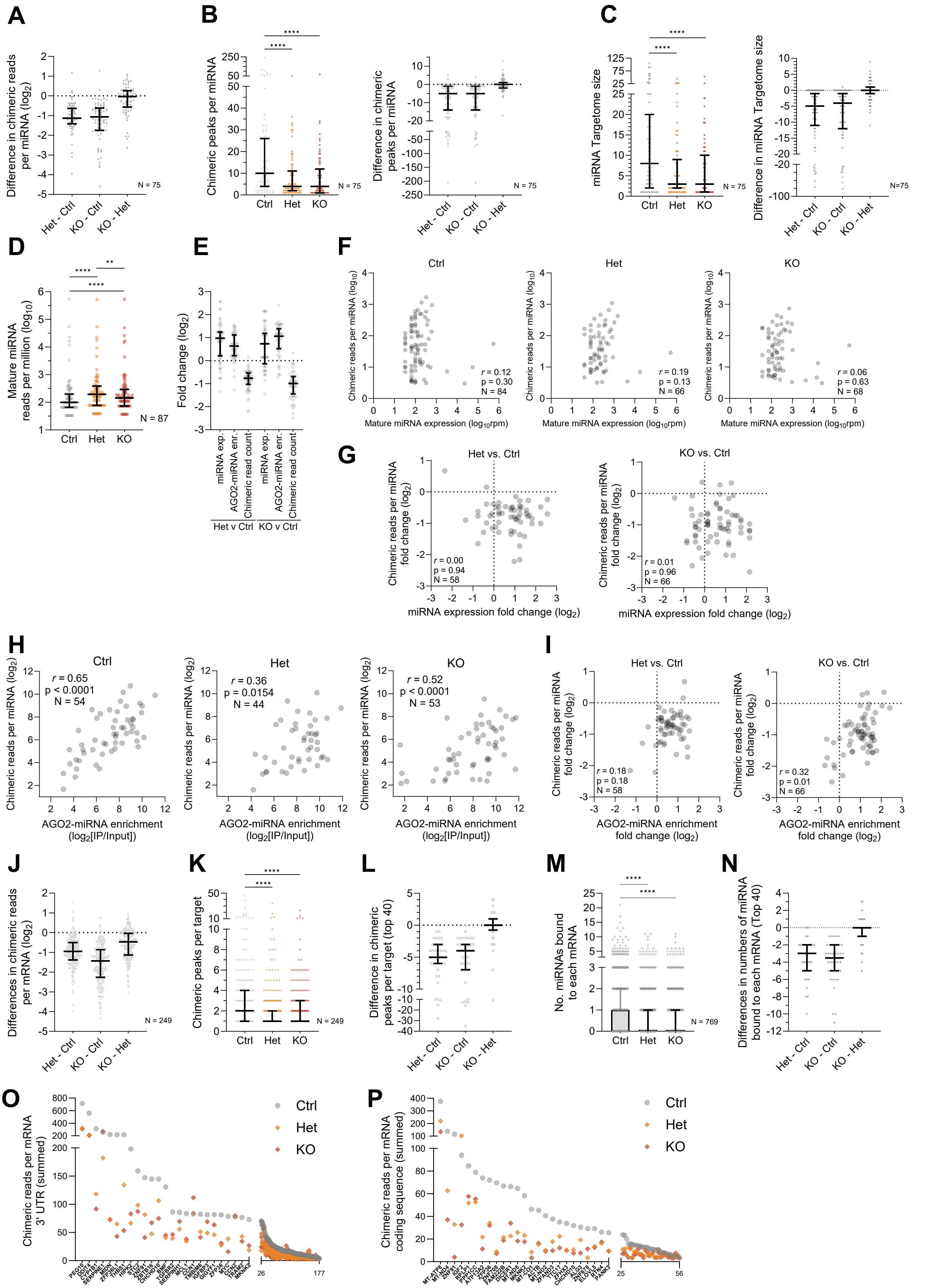

**Figure S5. LIMD1 deficiency constrains breadth and depth of targeting by individual AGO2–miRNAs**

**(A)** Differences in total chimeric reads ($\log_2$) per miRNA across 75 reproducibly detected miRNAs. **(B)** Distinct chimeric peaks per AGO2–miRNA, with paired differences (Wilcoxon matched-pairs signed-rank test; ****P<0.0001). **(C)** Targetome size per AGO2–miRNA, with paired differences, showing reduced targeting breadth per AGO2–miRNA in LIMD1-deficient cells (Wilcoxon test; ****P<0.0001). **(D)** General trend of increased mature miRNA expression per miRNA for miRNAs detected by chimeric-eCLIP (Wilcoxon test; **P<0.01, ****P<0.0001). **(E)** Fold change ($\log_2$) in miRNA expression, AGO2–miRNA enrichment, and chimeric reads, showing widespread loss of target binding despite increased expression and AGO2 loading. **(F)** Lack of correlation (Spearman) between miRNA expression ($\log_{10}$ RPM) and chimeric reads, indicating expression does not primarily drive targeting. **(G–I)** Correlation analyses of expression, AGO2 loading, and targeting. **(G)** No correlation between changes in expression and chimeric reads per miRNA in Het or KO versus Ctrl. **(H)** Positive correlation between AGO2 enrichment and chimeric reads, linking loading to targeting. **(I)** Positive correlation between AGO2 enrichment changes and chimeric reads, showing increased loading may partially offset targeting loss, though most miRNAs still show reduced reads. Overall, despite increased miRNA expression and AGO2 loading, chimeric reads declined. **(J)** Differences in total chimeric reads per target transcript. **(K, L)** Number of chimeric peaks per target transcript **(K)** and differences for the top 40 targets **(L)** (Wilcoxon test; ****P<0.0001). **(M, N)** Number of bound AGO2–miRNAs per target (M) and differences for the top 40 targets (N) (Wilcoxon test; ****P<0.0001). **(O, P)** Reduced chimeric reads per transcript at the **(O)** 3′UTR and **(P)** CDS, showing less AGO2–miRNA binding across both regions in LIMD1-deficient cells.

#### Figure S6. LIMD1 shapes AGO2–miRNA targeting landscapes by governing motif preferences and positional footprints

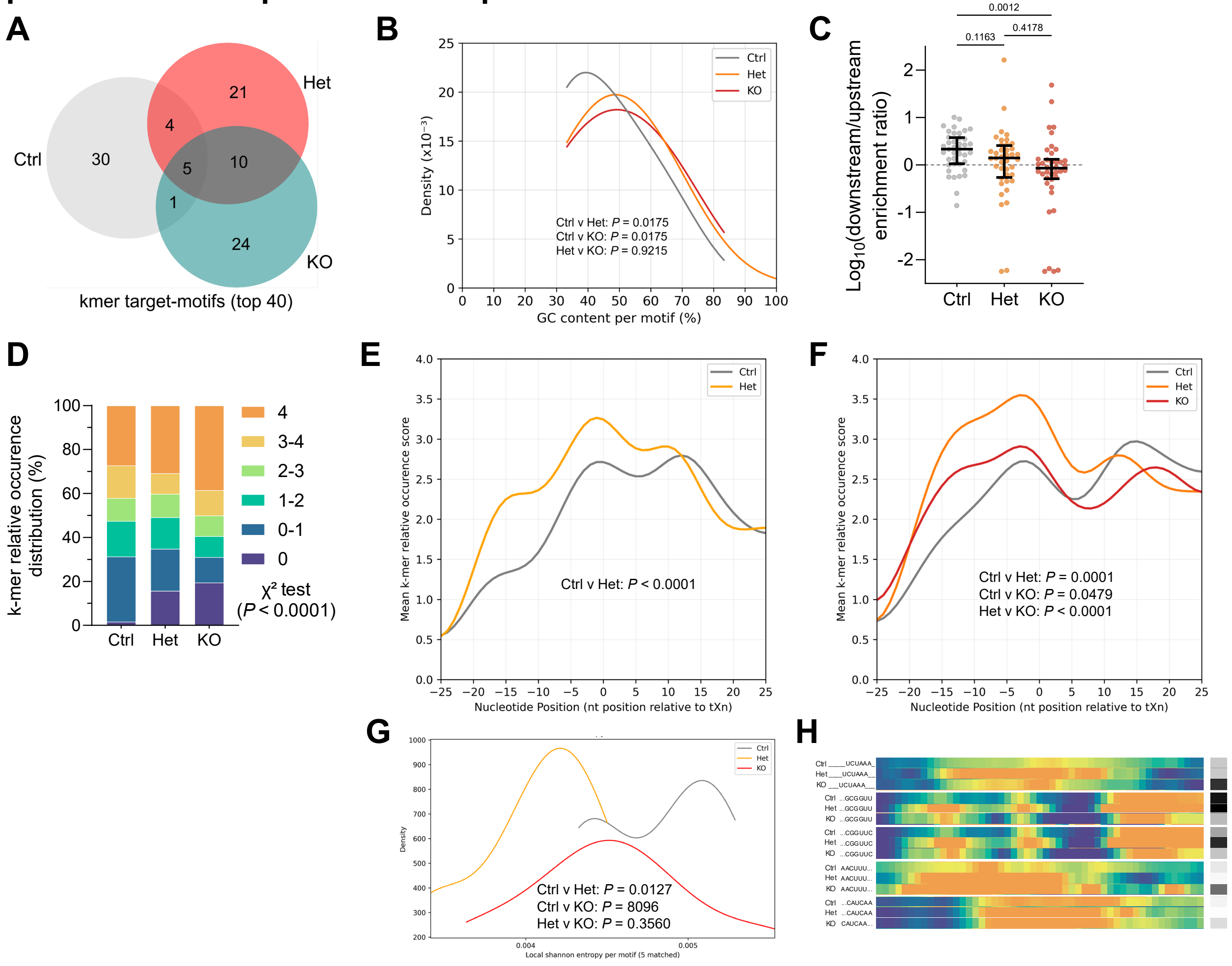


**Figure S6. LIMD1 shapes AGO2–miRNA targeting landscapes by governing motif preferences and positional footprints**

**(A)** Venn diagram of distinct and overlapping top 40 target motifs in Ctrl, Het, and KO. **(B)** Seed GC content of canonical interactions, showing a shift toward GC-rich motifs among interactions retained in LIMD1-deficient cells (kernel density, Gaussian σ = 3.33; Kruskal–Wallis with multiplicity correction). **(C)** Relative downstream/upstream enrichment ratio ($\log_2$) per motif among top 40 motifs; ratios >0 indicate downstream (3′) bias, <0 indicate upstream (5′) bias relative to tXn. **(D)** Distribution of RtXn values (0 = no binding, 1 = background, 4 = strong enrichment), with more zero-occurrence motifs and fewer with intermediate enrichment (1–<4) in LIMD1-deficient cells ($\chi^2$ test). **(E, F)** Motifs shared between Ctrl and Het (n = 9) **(E)** or Ctrl, Het, and KO (n = 5) **(F)** show an upstream (5′) shift in binding relative to tXn in LIMD1-deficient cells, even for identical motifs (Wilcoxon test). **(G, H)** Shannon entropy per motif for the five motifs common to all groups indicates that LIMD1 deficiency produces a sharper, punctate binding footprint even among identical target-motifs (corrected matched one-way ANOVA).

### Figure S7. LIMD1 deficiency de-represses miRNA targets in hSAECs and LUAD

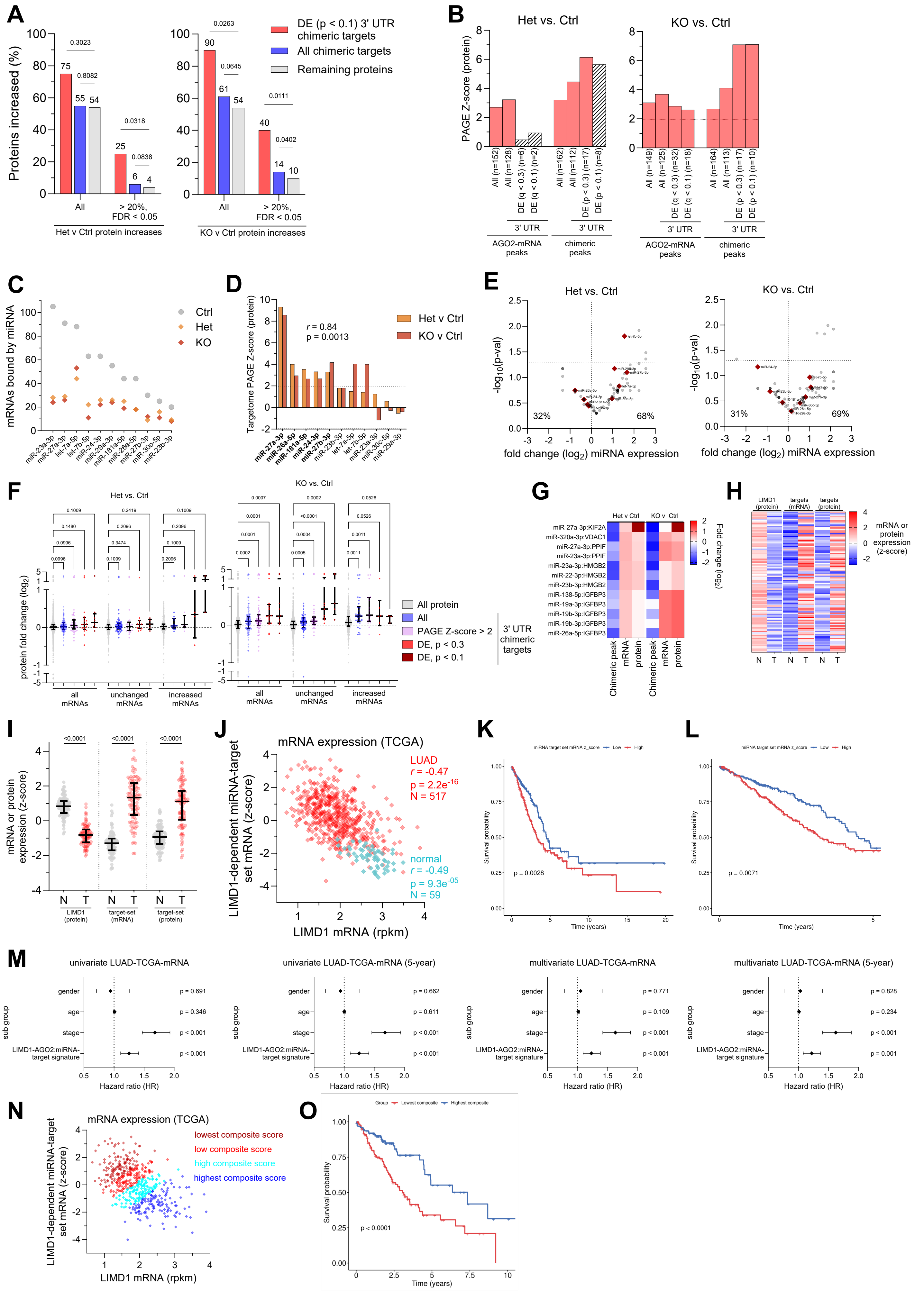

**Figure S7. LIMD1 deficiency de-represses miRNA targets in hSAECs and LUAD**

**(A)** Enrichment of all chimeric targets and top de-enriched ($P<0.1$) 3′UTR chimeric targets among increased proteins ($log_2FC>0$) and proteins increased >20% with FDR<0.05 (Fisher's exact test). **(B)** PAGE Z-scores showing significant protein increases for AGO2–miRNA targets in Het and KO versus Ctrl. Higher Z-scores were observed for chimeric versus AGO2–mRNA targets, 3′UTR versus all peaks, and top de-enriched 3′UTR peaks versus all 3′UTRs; $Z>1.96$ (dotted line) indicates significance ($P<0.05$). Dashed bars indicate insufficient protein numbers for PAGE analysis. **(C)** Fewer experimentally identified target mRNAs per miRNA in Het/KO versus Ctrl for miRNAs included in PAGE analysis. **(D)** Strong correlation of protein changes between Het and KO versus Ctrl indicates LIMD1-specific effects of target-dysregulation. **(E)** Expression changes of miRNAs with derepressed targetomes show several with increased abundance, confirming derepression is not due to reduced miRNA levels. **(F)** Protein fold changes ($log_2$) for 3′UTR chimeric targets (blue), targets of miRNAs with PAGE $Z>2$ (purple), and de-enriched chimeric targets ($P<0.1$, dark red), across all, unchanged (0.8–1.2-fold), and increased (>1.2-fold) mRNAs, showing elevated protein levels even without mRNA changes (Kruskal–Wallis with correction). **(G)** Chimeric peak, mRNA, and protein $log_2FC$ for the five genes in the target-signature set. **(H, I)** LIMD1 protein and target-set mRNA/protein z-scores in paired NAT (N) and tumour (T) samples (n=99) show reduced LIMD1 with increased target expression from NAT to tumour (corrected RM one-way ANOVA); **(H)** heatmap of matched samples and **(I)** boxplot. **(J)** Pearson correlation of LIMD1 mRNA with target-set mRNA z-scores in TCGA-LUAD normal and tumour samples. **(K, L)** Kaplan–Meier survival in TCGA-LUAD stratified by target-set mRNA signature (high vs low, top vs bottom half); high expression predicts poorer overall **(K)** and 5-year **(L)** survival (log-rank *P* values shown). **(M)** Forest plots of Cox regression (univariate and multivariate, overall and 5-year) showing tumour stage and high target-set mRNA levels predict poor survival outcomes. **(N, O)** LIMD1–target set composite score in LUAD tumours: **(N)** scatterplot of LIMD1 mRNA versus target-set expression coloured by composite quartiles (z[LIMD1] − z[target set]); **(O)** Kaplan–Meier survival for highest versus lowest composite score quartiles; low composite score predicts poorer survival (log-rank *P* value shown).

### Figure S8. LIMD1 augments evolutionarily young yet thermodynamically strong interactions

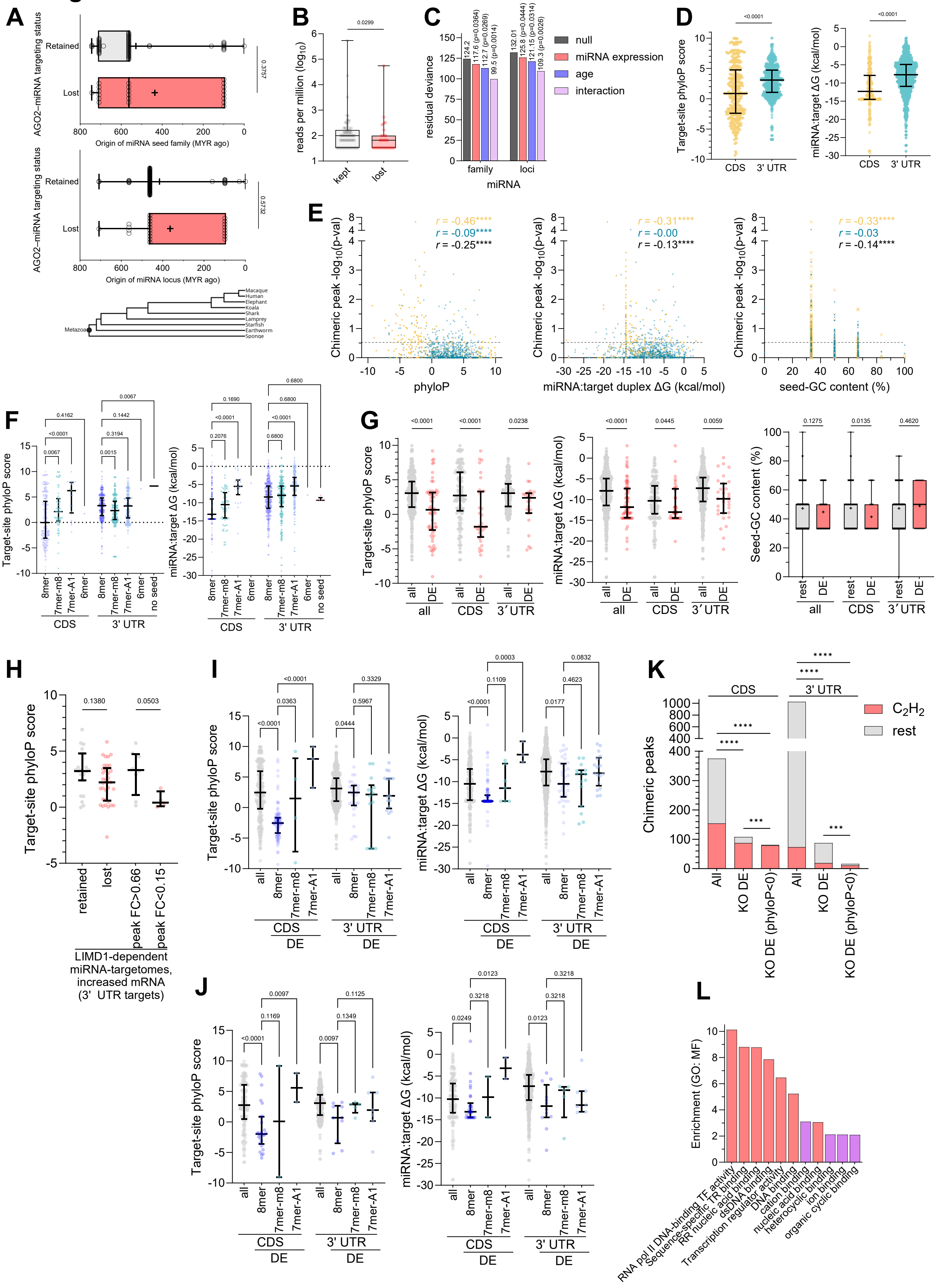

**Figure S8. LIMD1 augments evolutionarily young yet thermodynamically strong interactions**

**(A)** Evolutionary age (MYR) of miRNA seed families and loci categorized as retained or lost based on target engagement in LIMD1 KO versus Ctrl hSAECs. Younger miRNAs trended toward loss, but this did not approach significance (Kruskal–Wallis test). A phylogenetic tree provides context. **(B)** Lost miRNAs trend toward lower expression than retained miRNAs (Mann–Whitney test). **(C)** Logistic regression shows both expression level and evolutionary conservation independently predict miRNA retention. **(D)** Comparison of phyloP scores (>0 = increasing conservation, <0 = accelerated evolution) and free-binding energies between CDS and 3′UTR canonical target sites: CDS sites are less conserved but span a broader phyloP range and form more stable duplexes (more negative energies) (Mann–Whitney test). **(E)** Degree of LIMD1 dependence ($-\log_{10}$ p-value) versus features of canonical interactions. Spearman correlations are shown with phyloP conservation scores at target sites, predicted miRNA:target duplex stability ($\Delta G$, kcal/mol), and 6mer seed-GC content (%). CDS sites (yellow), 3′UTR sites (blue), and other regions (grey; 5′UTR, intron, miRNA, other) are indicated. Yellow, blue, and black r values denote correlations for CDS, 3′UTR, and all sites, respectively; significance indicated by asterisks (****$P<0.0001$). **(F)** Conservation and binding energies across site types (8mer, 7mer-m8, 7mer-A1) in CDS and 3′UTR regions: CDS 8mers are least conserved, CDS 7mer-A1 most conserved, and 8mers in both regions most stable (Kruskal–Wallis with correction). **(G)** For mRNAs with unchanged or increased abundance ($\log_2$FC>0), the most LIMD1-dependent canonical sites (significantly de-enriched in KO) show lower phyloP scores and more negative free-binding energies than the overall CDS or 3′UTR populations, excluding transcript-level effects (Kruskal–Wallis with correction). **(H)** For LIMD1-dependent miRNAs regulating 3′UTRs (miR-27a-3p, miR-27b-3p, miR-26a-5p, miR-24-3p, miR-181a-5p), sites with greatest chimeric read loss (FC<0.15) in transcripts of unchanged abundance are significantly less conserved than minimally affected sites (FC>0.66) (Kruskal–Wallis with correction). **(I, J)** Among all transcripts **(I)** or those unchanged/increased in abundance **(J)**, the most LIMD1-dependent canonical sites are enriched for poorly conserved, thermodynamically stable 8mers (Kruskal–Wallis with correction). **(K)** LIMD1-dependent sites (de-enriched [DE] in KO) are enriched for $C_2H_2$-ZNF genes, particularly in CDS; enrichment is even stronger for sites with phyloP<0 (accelerated evolution) ($\chi^2$ test; ***P<0.001, ****P<0.0001). **(L)** Gene Ontology (Molecular Function) analysis of LIMD1-dependent sites with phyloP<0 shows enrichment for DNA-binding and transcription functions (red) and small molecule/ion binding (purple) (FDR q<0.001; GOrilla, REViGO).

### Figure S9. Enlarged views of PEKA heatmaps from Fig. 5A–C

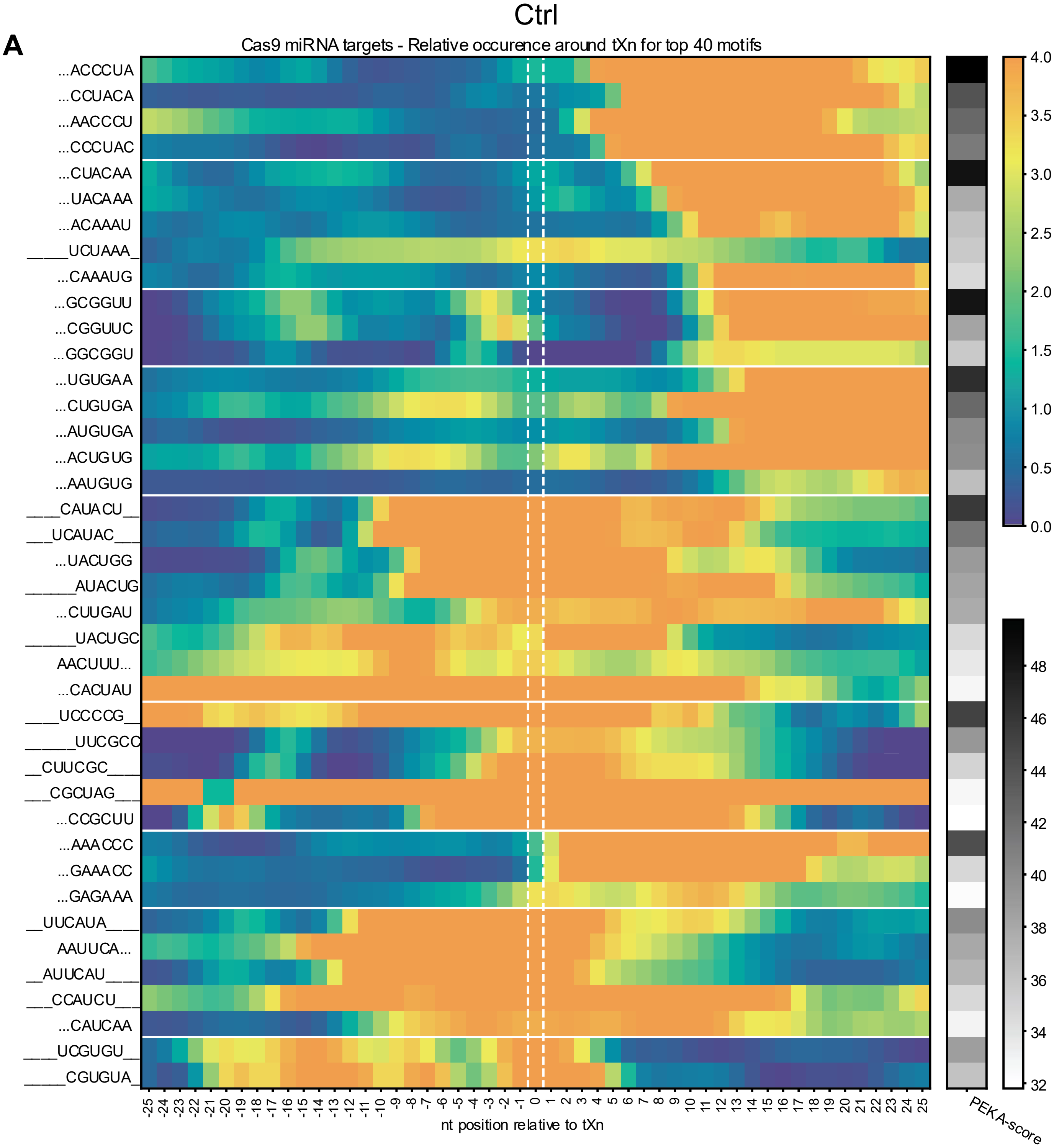


**Figure S9. Enlarged views of PEKA heatmaps from Fig. 5A–C**

**(A–C)** Enlarged versions of the (PEKA) heatmaps shown in Fig. 5A–C, corresponding to Ctrl (A), Het (B), and KO (C). Heatmaps depict motif relative occurrence (RtXn), defined as fold enrichment of k-mers near thresholded crosslink (tXn) sites relative to distal background. PEKA scores reflect enrichment strength and positional consistency. K-mers are clustered by sequence similarity and aligned by maximal RtXn; dotted lines indicate cases where maximal enrichment occurs >3 nt from the crosslink site. LIMD1 deficiency alters motif identity, positional distribution, and binding footprints.

### Figure S9. Enlarged views of PEKA heatmaps from Fig. 5A–C

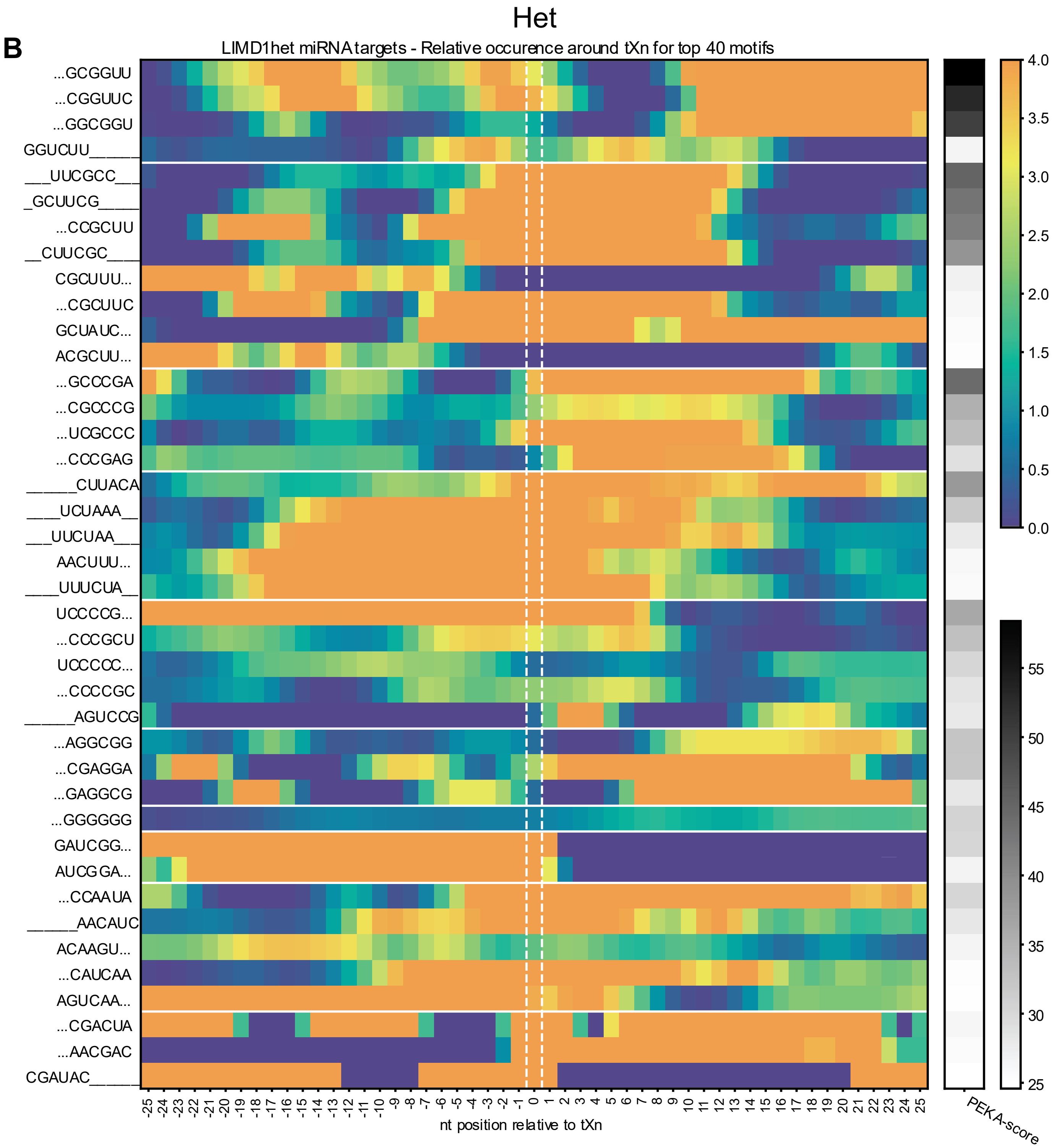

### Figure S9. Enlarged views of PEKA heatmaps from Fig. 5A–C

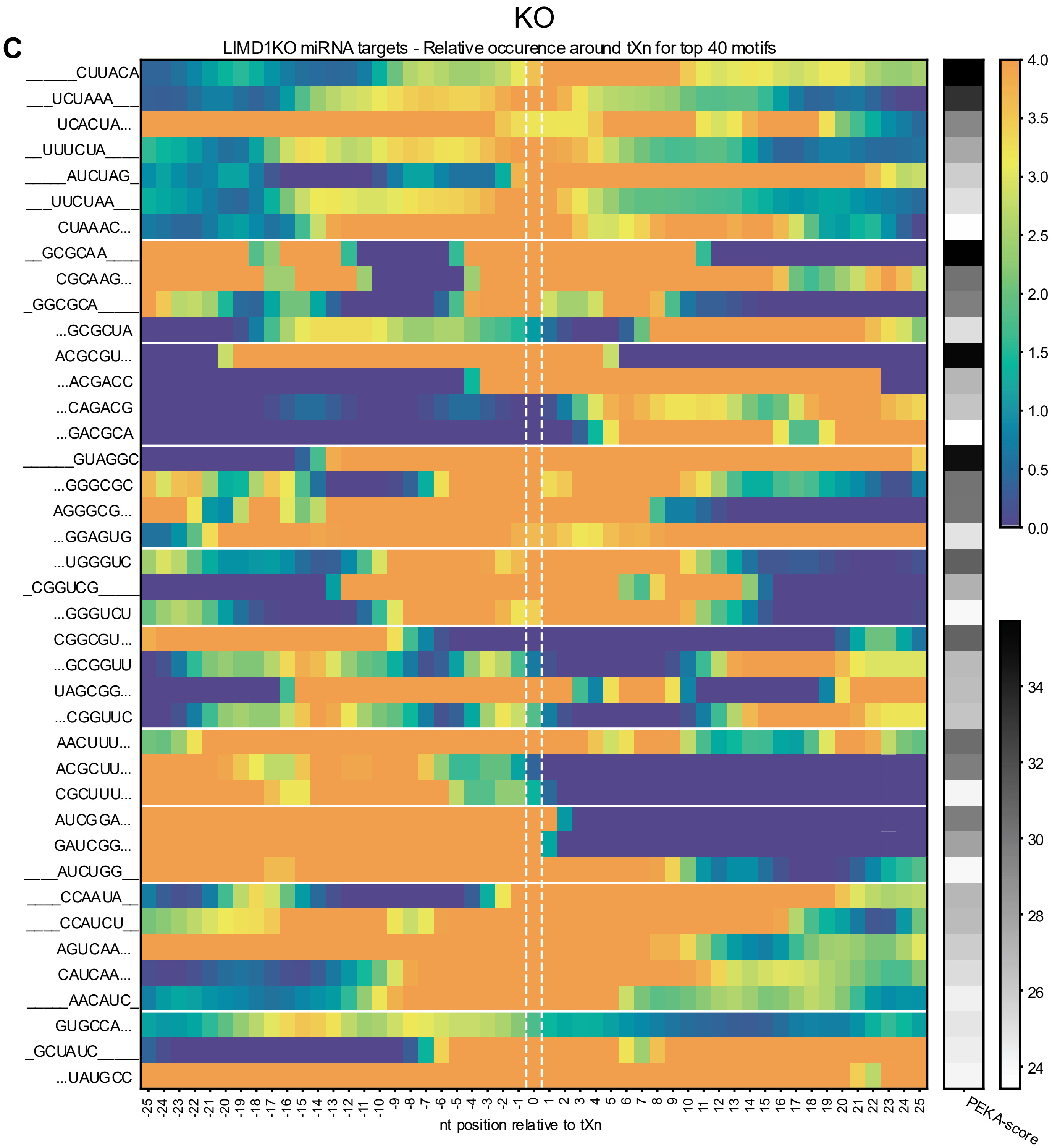

**Table S1. Summary of Sanger sequencing identifying editing events of LIMD1-targeted CRISPR-clones.**

| Sample | Editing event 1 | Editing Event 2 | Additional info? | Desired Genotype/FrameShift Indicated? |
|---|---|---|---|---|
| LIMD1 Het C6 | 1 deletion (T) after gDNA LIMD1 seq base 857 | No editing occurred (gDNA LIMD1 seq) | NA | Yes (*LIMD1*$^{+/-}$) |
| LIMD1 Het C8 | 5 base deletion (AAACT) at gDNA LIMD1 seq base 854. | No editing occurred (gDNA LIMD1 seq) | NA | Yes (*LIMD1*$^{+/-}$) |
| LIMD1 Het C11 | 3 bases before ATG start codon sequence stopped aligning | No editing occurred (gDNA LIMD1 seq) | Mis-alignment indicates large deletion | Yes (*LIMD1*$^{+/-}$) |
| LIMD1 KO C2 | 5 base deletion (AAACT) after LIMD1 gDNA base 853 | 5 base deletion (AAACT) after LIMD1 gDNA base 853 | C vs G at bp 900 indicates same editing event on two different alleles | Yes (*LIMD1*$^{-/-}$) |
| LIMD1 KO C10 | 1 insertion (T) after gDNA LIMD1 seq base 858 | 1 base deletion (T) after gDNA LIMD1 seq base 857 | NA | Yes (*LIMD1*$^{-/-}$) |
| LIMD1 KO C20 | 2 deletions (GC) at gDNA LIMD1 base 859. | 2 deletions (CT) at gDNA LIMD1 base 857. | NA | Yes (*LIMD1*$^{-/-}$) |

**Table S2. Summary of whole genome sequencing (WGS) results probing on-target CRISPR-Cas9-mediated editing events.**
Alignment E-values signify number of expected alignments of equivalent or greater score, reflecting strong sequence homology to the guide. VAF = variant allele frequency.

| Genotype and clone ID | On-target edit (WGS) | Alignment E-value | WGS evidence | Sanger phasing details |
|---|---|---|---|---|
| LIMD1 Het C6 | 1 bp del (CT→C) at Chr 3:45,595,010 | $4 \times 10^{-5}$ | 30× depth; VAF ~52% | Confirmed |
| LIMD1 Het C8 | 1 bp del (CT→C) at Chr 3:45,595,010 | $4 \times 10^{-5}$ | 30× depth; VAF ~48% | Confirmed |
| LIMD1 Het C11 | 4 bp del (TCGCA→T) at Chr 3:45,595,011 | $2 \times 10^{-5}$ | 33× depth; VAF ~50% | Confirmed |
| LIMD1 KO C2 | 5 bp del (GAAACT→G) at Chr 3:45,595,006 | $9 \times 10^{-6}$ | 32× depth; VAF > 95% | Confirmed |
| LIMD1 KO C10 | Exon-scale deletion (~2 kb) | n/a | ≥ 10 split reads & discordant pairs; ~50 % coverage drop | Compound het: +T ins after base 858; –T del after base 857 |
| LIMD1 KO C20 | Exon-scale deletion (~2 kb) | n/a | ≥ 10 split reads & discordant pairs; ~45% coverage drop | Biallelic 2 bp dels (ΔGC at 859; ΔCT at 857) |

**Table S3. Summary of Whole genome sequencing results probing putative off-target CRISPR-Cas9 editing events.**
Homology E-values signify number of expected alignments of equivalent or greater score, here reflecting weak sequence homology to the guide.

| Genotype and clone ID | Putative off-target edit? | Chromosome | Genomic Coordinates (GRCh38, bp) | Deletion length (bp) | Homology E-value | Genomic context |
|---|---|---|---|---|---|---|
| LIMD1 Het C6 | Yes | 2 | Chr 2:62,730,764–62,730,886 | 122 | 11 | Noncoding |
| LIMD1 Het C8 | No | – | – | – | – | – |
| LIMD1 Het C11 | No | – | – | – | – | – |
| LIMD1 KO C2 | No | – | – | – | – | – |
| LIMD1 KO C10 | Yes | 2 | Chr 2:62,730,772–62,730,879 | 107 | 9.7 | Noncoding |
| LIMD1 KO C20 | Yes | 2 | Chr 2:62,730,764–62,730,853 | 89 | 8.6 | Noncoding |

#### Table S4.

| qc_metrics | | | | | | | | | | | | |
|---|---|---|---|---|---|---|---|---|---|---|---|---|
| Sample | IP or Input | Initial reads | % pass trim | % rep elements | % uniquely aligned to genome | % PCR duplicates | Final nonchimeric reads | AGO2 clusters | AGO2 peaks | Final chimeric reads | % chimeras | Chimeric clusters |
| input_WT_R1_001 | Input | 39,844,858 | 92.03% | 64.80% | 33.65% | 22.18% | 3,379,282 | | | | | |
| input_Ctrl_C3_R1_001 | Input | 65,427,680 | 99.10% | 67.60% | 43.77% | 18.81% | 7,465,573 | | | | | |
| input_Ctrl_C4_tr2_R1_001 | Input | 74,443,442 | 97.87% | 66.91% | 40.41% | 21.47% | 7,650,401 | | | | | |
| input_Ctrl_C2_R1_001 | Input | 39,573,542 | 87.10% | 73.66% | 21.62% | 21.98% | 1,531,132 | | | | | |
| input_Ctrl_C4_tr1_R1_001 | Input | 44,723,159 | 97.55% | 63.75% | 34.40% | 20.18% | 4,343,195 | | | | | |
| input_LIMD1_Het_C6_tr1_R1_001 | Input | 35,619,172 | 75.60% | 64.25% | 29.30% | 19.65% | 2,266,277 | | | | | |
| input_LIMD1_Het_C8_R1_001 | Input | 46,253,733 | 88.26% | 72.28% | 23.02% | 22.75% | 2,011,613 | | | | | |
| input_LIMD1_Het_C11_R1_001 | Input | 72,187,939 | 94.02% | 60.33% | 30.83% | 23.48% | 6,350,342 | | | | | |
| input_LIMD1_Het_C6_tr2_R1_001 | Input | 68,990,119 | 99.37% | 62.19% | 36.21% | 20.96% | 7,418,744 | | | | | |
| input_LIMD1_KO_C10_R1_001 | Input | 43,115,873 | 90.42% | 51.59% | 40.98% | 20.43% | 6,154,821 | | | | | |
| input_LIMD1_KO_C2_tr1_R1_001 | Input | 38,989,398 | 87.07% | 68.23% | 27.52% | 21.62% | 2,325,908 | | | | | |
| input_LIMD1_KO_C20_R1_001 | Input | 66,900,528 | 98.23% | 69.19% | 35.90% | 20.18% | 5,802,502 | | | | | |
| input_LIMD1_KO_C2_tr2_R1_001 | Input | 63,084,068 | 99.05% | 64.02% | 43.74% | 17.93% | 8,070,176 | | | | | |
| IP_WT_R1_001 | IP | 43,185,429 | 93.18% | 35.61% | 38.00% | 24.82% | 7,402,362 | 203,241 | 5,127 | 68,147 | 0.91% | 1,027 |
| IP_Ctrl_C3_R1_001 | IP | 74,775,141 | 98.43% | 38.13% | 28.75% | 17.92% | 10,745,618 | 568,912 | 8,079 | 87,380 | 0.81% | 1,331 |
| IP_Ctrl_C4_tr2_R1_001 | IP | 91,933,843 | 98.72% | 36.13% | 29.86% | 21.59% | 13,571,213 | 568,933 | 7,409 | 82,827 | 0.61% | 1,476 |
| IP_Ctrl_C2_R1_001 | IP | 62,766,609 | 97.45% | 52.35% | 27.23% | 23.72% | 6,052,533 | 568,924 | 3,686 | 67,515 | 1.10% | 1,476 |
| IP_Ctrl_C4_tr1_R1_001 | IP | 54,872,857 | 98.01% | 40.59% | 43.18% | 21.71% | 10,801,829 | 568,962 | 8,729 | 122,848 | 1.12% | 2,050 |
| IP_LIMD1_Het_C6_tr1_R1_001 | IP | 58,207,062 | 96.29% | 49.19% | 32.77% | 22.05% | 7,273,949 | 533,691 | 4,679 | 52,724 | 0.72% | 784 |
| IP_LIMD1_Het_C8_R1_001 | IP | 61,930,112 | 87.28% | 51.63% | 27.64% | 21.26% | 5,691,103 | 533,699 | 2,515 | 29,161 | 0.51% | 526 |
| IP_LIMD1_Het_C11_R1_001 | IP | 81,083,849 | 98.28% | 46.45% | 26.65% | 22.15% | 8,852,616 | 533,680 | 4,347 | 41,289 | 0.46% | 665 |
| IP_LIMD1_Het_C6_tr2_R1_001 | IP | 82,135,404 | 97.84% | 38.33% | 32.46% | 20.85% | 12,730,630 | 533,702 | 5,395 | 69,155 | 0.54% | 979 |
| IP_LIMD1_KO_C10_R1_001 | IP | 45,212,401 | 92.73% | 35.69% | 34.53% | 21.10% | 7,346,652 | 530,767 | 4,715 | 33,595 | 0.46% | 547 |
| IP_LIMD1_KO_C2_tr1_R1_001 | IP | 47,421,688 | 93.44% | 55.42% | 28.17% | 20.64% | 4,416,552 | 530,771 | 3,985 | 34,583 | 0.78% | 652 |
| IP_LIMD1_KO_C20_R1_001 | IP | 74,234,778 | 98.13% | 46.76% | 26.30% | 20.49% | 8,111,229 | 530,808 | 4,948 | 51,718 | 0.63% | 758 |
| IP_LIMD1_KO_C2_tr2_R1_001 | IP | 82,288,388 | 98.32% | 36.42% | 28.09% | 19.94% | 11,567,691 | 530,794 | 4,976 | 51,165 | 0.44% | 703 |

#### Table S5.

| qc_metrics (summary) | | | | | | | | | | | | |
|---|---|---|---|---|---|---|---|---|---|---|---|---|
| Sample | IP or Input | Initial reads | % pass trim | % rep elements | % uniquely aligned to genome | % PCR duplicates | Final nonchimeric reads | AGO2 clusters | AGO2 peaks | Final chimeric reads | % chimeras | Chimeric clusters |
| input_Ctrl | Input | 56,041,956 | 95.41% | 67.98% | 35.05% | 20.61% | 5,247,575 | | | | | |
| input_LIMD1_Het | Input | 55,762,741 | 89.31% | 64.76% | 29.84% | 21.71% | 4,511,744 | | | | | |
| input_LIMD1_KO | Input | 53,022,467 | 93.69% | 63.26% | 37.04% | 20.04% | 5,588,352 | | | | | |
| IP_Ctrl | Input | 71,087,113 | 98.15% | 41.80% | 32.26% | 21.24% | 10,292,798 | 568,933 | 6,976 | 90,143 | 0.91% | 1,583 |
| IP_LIMD1_Het | Input | 70,839,107 | 94.92% | 46.40% | 29.88% | 21.58% | 8,637,075 | 533,693 | 4,234 | 48,082 | 0.56% | 739 |
| IP_LIMD1_KO | Input | 62,289,314 | 95.66% | 43.57% | 29.27% | 20.54% | 7,860,531 | 530,785 | 4,656 | 42,765 | 0.58% | 665 |

**Table S4-5. Quality control metrics of chimeric-eCLIP experiment**

**4)** This table summarizes sequencing and processing statistics for all chimeric-eCLIP libraries (two biological replicates per genotype) and their matched inputs. For each sample, we report:

IP or input: immunoprecipitation (IP) versus input control.
Initial reads: total raw reads obtained.
% pass trim: fraction of reads retained after adapter and quality trimming.
% rep elements: fraction of reads mapping to repetitive elements.
% uniquely aligned to genome: fraction of non-repetitive reads that map uniquely to the reference genome.
% PCR duplicates: fraction of uniquely aligned reads flagged as PCR duplicates.
Final non-chimeric reads: number of unique, non-chimeric reads used for peak calling.
AGO2 clusters: number of enriched binding clusters called genome-wide.
AGO2 peaks: number of reproducible peaks after irreproducible discovery rate (IDR) filtering. A peak is defined as a cluster with log2 fold enrichment > 3 and p-value < 0.001.
Final chimeric reads: number of reads containing chimeric (miRNA–mRNA) junctions.
% chimeras: proportion of final non-chimeric reads that yielded chimeric alignments.
Chimeric clusters: number of unique miRNA–target interaction sites identified.

These metrics confirm high library complexity, efficient removal of artifacts, and robust identification of AGO2–miRNA crosslinked sites across all genotypes, as well as consistency of differences between sample groups. One replicate was performed on WT cells (non-CRISPR-targeted and no single-cell selection) hSAECs for benchmarking.

**5)** Summary (means) of above per input or IP sample for each sample group.
